# Dynamics of sex-biased gene expression during development in the stick insect *Timema californicum*

**DOI:** 10.1101/2021.01.23.427895

**Authors:** Djordjevic Jelisaveta, Dumas Zoé, Robinson-Rechavi Marc, Schwander Tanja, Parker Darren James

**Affiliations:** Department of Ecology and Evolution, University of Lausanne, Lausanne, Switzerland; Swiss Institute of Bioinformatics, Lausanne, Switzerland

**Keywords:** Sex-biased gene expression, development, *Timema* stick insects, *Drosophila*, sexual dimorphism

## Abstract

Sexually dimorphic phenotypes are thought to arise primarily from sex-biased gene expression during development. Major changes in developmental strategies, such as the shift from hemimetabolous to holometabolous development, are therefore expected to have profound consequences for the dynamics of sex-biased gene expression. However, no studies have previously examined sex-biased gene expression during development in hemimetabolous insects, precluding comparisons between developmental strategies. Here we characterized sex-biased gene expression at three developmental stages in a hemimetabolous stick insect (*Timema californicum*): hatchlings, juveniles, and adults. As expected, the proportion of sex-biased genes gradually increased during development, mirroring the gradual increase of phenotypic sexual dimorphism. Sex-biased genes identified at early developmental stages were generally consistently male- or female-biased at later stages, suggesting their importance in sexual differentiation. Additionally, we compared the dynamics of sex-biased gene expression during development in *T. californicum* to those of the holometabolous fly *Drosophila melanogaster* by reanalyzing publicly available RNA-seq data from third instar larval, pupal and adult stages. In *D. melanogaster*, 84% of genes were sex-biased at the adult stage (compared to only 20% in *T. californicum*), and sex-biased gene expression increased abruptly at the adult stage when morphological sexual dimorphism is manifested. Our findings are consistent with the prediction that the dynamics of sex-biased gene expression during development differ extensively between holometabolous and hemimetabolous insect species.

## Introduction

Males and females often have divergent evolutionary interests, resulting in sex-specific selection pressures and ultimately the evolution of sexually dimorphic phenotypes (Lande 1980; Khila *et al*. 2012). Studies investigating how sexually dimorphic phenotypes are generated have shown the importance of differential gene expression between the sexes, suggesting that sex-specific selection is the major driving force behind the evolution of sex-biased gene expression (Mank 2017). The relationship between sexual dimorphism and sex-biased gene expression has been largely studied at the adult stage when sexual dimorphism is completely manifested and reproductive interests between males and females are most different (Mank 2017). However, sex-biased gene expression has also been found at early developmental stages of many species, well before any phenotypic sexual dimorphism becomes apparent (Lowe *et al*. 2015; Paris *et al*. 2015). This suggests that expression patterns in early developmental stages are also under sex-specific selection pressures (Mank *et al*. 2010; Hale *et al*. 2011; Zhao *et al*. 2011; Perry *et al*. 2014; Ingleby *et al*. 2015).

Because we lack detailed studies of sex-biased gene expression during development, we do not know how consistent or dynamic the expression of the sex-biased genes is, nor how these dynamics relate to changes in phenotypic sexual dimorphism. For example, although we expect to see an overall increase in sex-biased gene expression during development (Ingleby *et al*. 2015; Mank 2017), it is not clear whether this results from the gradual increase of sex-bias of a set of genes or if sex-biased genes at early stages are largely different to those at later stages (Mank 2017). Furthermore, sex-biased gene expression can be considered to reflect a broad measure of sexual dimorphism, including physiological and behavioral traits, and may therefore be more representative than dimorphism quantified using external morphology. However, whether the extent of sexual dimorphism and of sex-biased gene expression are generally correlated remains poorly known.

In insects there are two major developmental strategies, and developmental patterns of sex-biased gene expression could differ between them. Holometabolous insects have morphologically and ecologically distinct larval, pupal, and adult stages, and phenotypic sexual dimorphism is commonly prominent only at the adult stage. Contrastingly, hemimetabolous insects go through gradual morphological changes and sexual differentiation, and the nymphal stages morphologically resemble adults (Chen *et al*. 2010). These distinct dynamics for sexual morphological differentiation are expected to be mirrored by sex-biased gene expression, with abrupt versus gradual increases in the proportion of sex-biased genes during development. However, studies of sex-biased gene expression during development have thus far only been conducted in holometabolous insects (Magnusson *et al*. 2011; Rago *et al*. 2020), with the main focus on *Drosophila* (Perry *et al*. 2014; Ingleby *et al*. 2015), precluding comparisons between species with different developmental strategies. For example, more than 50% of the transcriptome is sex-biased in pre-gonad tissue in juvenile stages of *Drosophila melanogaster* (Perry *et al*. 2014) but similar data are not available for hemimetabolous species. Furthermore, sex-biased genes in reproductive tissues provide only partial insight into total sexual dimorphism, since it excludes differences beyond sexual organs, such as secondary sexual traits.

Here, we studied the dynamics of sex-biased gene expression across development in a hemimetabolous insect, *Timema californicum. Timema* are sexually dimorphic walking-stick insects, found in California (Vickery and Sandoval 2001). Males are smaller than females, with different coloration, and have conspicuous cerci used for holding on to the female during copulation. We selected three developmental stages (Fig. 1a): the hatchling stage where sexes are phenotypically identical, the third juvenile stage with minor phenotypic differences, and the adult stage, with pronounced sexual dimorphism. We investigated whether sex-biased genes are recruited in a stage-specific manner, and if the magnitude of sex-bias increases with development. We further explored sequence evolution rates of sex-biased genes in pre-adult stages, to determine if they show elevated rates as commonly observed for sex-biased genes at adult stages in many species (Ellegren and Parsch 2007; Grath and Parsch 2012; Perry *et al*. 2014).

**Fig. 1.**
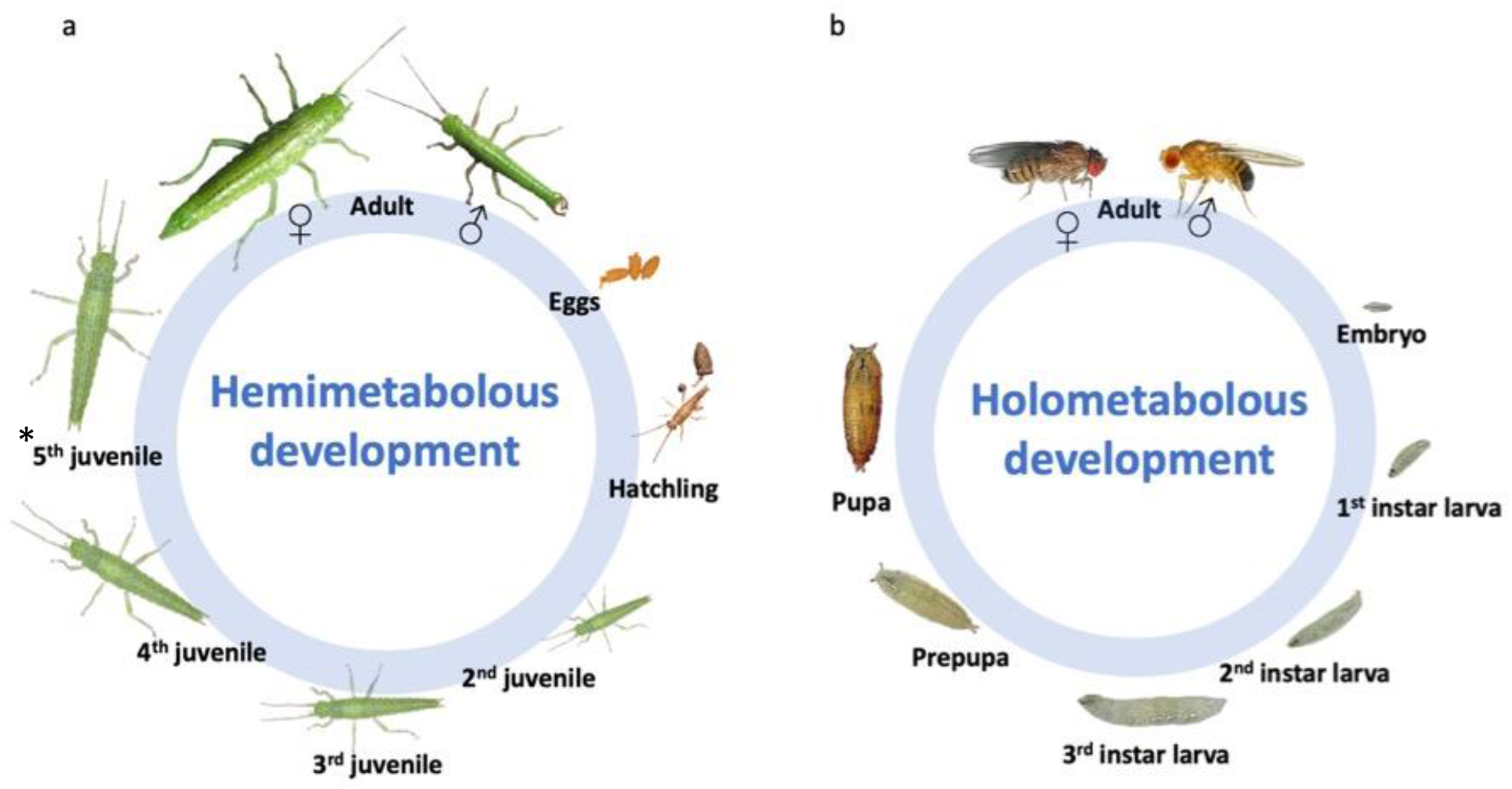
Life cycles of (a) the stick insect *T. californicum* and (b) the fly *D. melanogaster*. *represents a stage present only in females, as *Timema* males have one moult fewer than females. Hatchling and egg photographs of *Timema* were kindly provided by Bart Zijlstra (http://www.bartzijlstra.com), juvenile and adult stages by Jelisaveta Djordjevic. Life cycle of *D. melanogaster* modified from Weigmann *et al*. (2003).

We then compared the dynamics of sex-biased gene expression during development in the hemimetabolous stick insect to those of the holometabolous fly *Drosophila melanogaster* by reanalyzing publicly available RNA-seq data from 3rd. instar larvae, pupal and adult stages, using the same pipeline as for *T. californicum*. In *D. melanogaster*, both larval and pupal stages have little morphological sexual dimorphism, while the adult stage has extensive dimorphism (Fig. 1b). We hypothesized that sex biased gene expression follows the establishment of morphological sexual dimorphism, with a gradual increase in sex-biased gene expression during development in *T. californicum*, and an abrupt increase at the adult stage in *D. melanogaster*.

We showed that the proportion of sex-biased genes gradually increases during development in *T. californicum*, mirroring the gradual differentiation of phenotypic sexual dimorphism in hemimetabolous insects. Sex-biased genes from early developmental stages largely remained sex-biased at later stages. Finally, our findings are consistent with the prediction that the dynamics of sex-biased gene expression during development differs extensively between holometabolous and hemimetabolous insect species.

## Methods

### Sample collection and preservation

*T. californicum* eggs hatch in early winter. Upon hatching, insects moult several times before reaching maturity in late spring (Sandoval 1994). Developmental stages (especially adults and hatchlings) do not co-occur temporally, hence samples from different stages cannot be collected simultaneously. Adults (4 males, 4 females) and juveniles (4 males, 3 females) were collected in California (Saratoga County) in 2015 and 2016, respectively. They were fed with artificial medium for two days to prevent contamination with plant cells from the gut, and subsequently frozen at −80°C. Hatchlings were obtained from the eggs laid by field-collected adults in 2015. Hatchlings were flash frozen and stored at −80 °C. Sexes cannot be distinguished morphologically at the hatchling stage, thus single individuals were extracted to allow for later identification of sex via genotyping (see below) and 5 male and 4 female hatchlings were used.

### RNA extraction and sequencing

Individuals (whole-bodies) from all developmental stages were mechanically homogenized with beads (Sigmund Linder) in liquid nitrogen. We then added 900 ul Trizol (Life Technologies), followed by 180 ul chloroform and 350 ul ethanol. The aqueous layer was then transferred to RNeasy MinElute Columns (Qiagen). Upon RNA extraction, samples were treated with DNase Turbo Kit (Life Tech) following the manufacturer’s protocol. Total RNA from the hatchlings was extracted with MagMaxTM Express Robot (AB Applied Biosystems) using a MagMAXTM-96 Total RNA Isolation Kit from Ambion (Life Technologies) following the manufacturer’s protocol, but without DNAse at this step to preserve DNA for the sex determination of hatchlings via genotyping (see below). Following RNA extraction, hatchling samples were treated with DNase Turbo Kit (Life Tech) following the manufacturer’s protocol. The quality and quantity of the extracted RNA was then measured using a NanoDrop and Bioanalyzer. Library preparations (one for each individual) were done using the Illumina TruSeq Stranded Total RNA kit, upon which samples were sequenced together in six lanes. Paired-end sequencing with a read length of 100 bp was done on a HiSeq2000 platform at the GTF (Genomic Technologies Facility, Centre of Integrative Genomics, Lausanne, Switzerland).

### Hatchling sex identification

To identify the sex of *T. californicum* hatchlings, we developed microsatellite markers on the *T. californicum* X chromosome scaffolds reported by Parker *et al*. 2021. *Timema* have an XX/X0 sex determination system (Schwander and Crespi 2009) meaning X-linked regions will be present in two copies in females and one copy in males. Consequently, females can feature heterozygous genotypes for polymorphic X-linked markers, while males are invariably hemizygous. Candidate microsatellite markers were designed with msatcommander (v. 1.08, default options) (Faircloth 2008). A selection of four candidate markers were tested for polymorphism and sex-linkage using 15 adult males and 15 adult females in two Multiplex PCR Kit reactions (Qiagen; see Supplemental Table 1 for primer sequences and Supplemental Table 2 for PCR conditions). Microsatellite alleles were then determined with an ABI3100 machine (Applied Biosystem) and Genemapper v.4.1 (Applied Biosystems 2009). Across microsatellite markers, all 15 tested females were heterozygous at minimum two markers, whereas males were invariably hemizygous at all markers, confirming the predictions for *Timema* X-linked markers (Supplemental Table 2). We thus genotyped the nine hatchlings at the four microsatellite markers to identify their sex, and were able to identify four females and five males that were used for the transcriptome study (Supplemental Table 2).

### Raw data quality control, mapping and read counting

The quality of the reads was checked using FastQC v.0.11.2 (Andrews 2014). We used Cutadapt v. 2.3 with Python v. 3.5.2 (Martin 2011) to remove adapter sequences. Low quality bases at both ends of the reads were trimmed when below a Phred score of 10. Bases with an average Phred score below 20 in a 4bp sliding window were trimmed. Finally, reads with a length below 80 bp were removed with Trimmomatic v. 0.36 (Bolger *et al*. 2014). Trimmed reads were then mapped to the reference *T. californicum* genome from Jaron *et al*. (2020) using STAR v. 2.6.0c (Dobin *et al*. 2013). HTSeq v.0.9.1 (Anders *et al*. 2014) was used to count the number of reads uniquely mapped to each gene, with the following options (htseq-count -f bam -r name -s reverse -t gene -i ID -m union --nonunique none).

### *Drosophila melanogaster* data

*D. melanogaster* RNA-seq whole body data from Ingleby *et al*. (2016) was downloaded from SRA (accession number SRP068235). To mirror the data available for *Timema* and to facilitate comparisons, four replicates per sex and three developmental stages from different hemiclonal lines were used. A complete list of samples is provided in the supplementary materials (Supplemental Table 3). Although reads were deposited as paired-end, we found that most of them were unpaired, thus we used only the forward reads for analysis. Reads were trimmed, mapped to the reference genome (FlyBase, r6.23) (Gramates *et al*. 2017) and counted using the methods described above.

### Differential gene expression analysis

Differential expression between the sexes was performed using edgeR v.3.16.5 (Robinson *et al*. 2009; McCarthy *et al*. 2012) in RStudio v.1.2.5019 (Team 2019). This analysis fits a generalized linear model following a negative binomial distribution to the expression data and calculates a p-value associated with the hypothesis that gene counts are similar between experimental groups. EdgeR implements a normalization method (trimmed mean of M values, TMM) to account for composition bias. We analyzed data separately for each developmental stage. We filtered out genes with low counts; we required a gene to be expressed in a majority of male or female libraries dependent on the number of replicates per sex (i.e. a minimum three libraries when replicate number is >= 4, two libraries when replicate number is = 3), with expression level > 0.5 CPM (counts per million). To test for differential gene expression between the sexes, we used a generalized linear model with a quasi-likelihood F-test, with contrasts for each stage between females and males (Chen *et al*. 2016). To correct for multiple testing, we applied the Benjamini-Hochberg method (Benjamini and Hochberg 1995) with an alpha of 5%. Genes with significantly higher expression in males were considered as male-biased, genes with significantly higher expression in females as female-biased, and the genes without significant differential expression between sexes as un-biased. To identify genes showing a significant sex by developmental stage interaction we used a similar GLM modelling approach but with all stages included. We used SuperExactTest v.0.99.4 (Wang *et al*. 2015) to test if the overlap of sex-biased genes between the three developmental stages was greater than what is expected by chance. To visualize the overlap of sex-biased genes between developmental stages we used VennDiagram v.1.6.20 (Chen and Boutros 2011). We visualized the log_2_FC sex-biased gene expression across developmental stages, using pheatmap v.1.0.12 (Kolde 2018). We tested the Spearman’s correlation of sex-bias (log_2_FC) between the developmental stages, using cor.test from the R package stats v 4.1.1. To describe the increases in the number of sex-biased genes during development, we calculated the effect sizes for each of the two proportions of neighboring developmental stages, using the pwr v. 1.3-0 (Cohen 1988).

### Stage specific gene expression

Median gene expression (CPM) values were calculated for each developmental stage and sex. These values were used to calculate Tau, an index of gene expression specificity during development (Yanai *et al*. 2004; Liu and Robinson-Rechavi 2018). Values of Tau range from zero (broadly expressed during development) to one (gene expressed in only one stage). Tau is generally used to quantify the tissue-specificity of gene expression (Yanai *et al*. 2004). Here we applied the same principle and formula to calculate stage specificity. To visualize the results, we made boxplots using ggplot2 v. 3.3.2 (Wickham 2016).

### Divergence rates

We calculated values of sequence divergence rates (dN/dS) along the branch leading to *T. californicum* after the split with *T. poppensis* as described in (Jaron *et al*. 2020). Briefly, branch-site models with rate variation at the DNA level (Davydov *et al*. 2019) were run using the Godon software (https://bitbucket.org/Davydov/godon/, version 2020-02-17, option BSG --ncat 4) for each gene with an ortholog found in at least 6 species of *Timema* (including *T. poppensis*). Godon estimates the proportion of sites evolving under purifying selection (p0), neutrality (p1), and positive selection. We used only sites evolving under purifying selection or neutrality to calculate dN/dS. To test for differences in sequence divergence rates (dN/dS) between different gene categories (female-biased, male-biased and un-biased), we used Wilcoxon tests (ggpubr v.0.2.5, in R) with p-values adjusted for multiple comparisons using the Benjamini-Hochberg method (Haynes 2013). Note that some of the *T. californicum* sex-biased genes had no identified ortholog in other *Timema* species, thus we do not have sequence evolution rates for all sex-biased genes.

Furthermore, we used a partial correlation between the strength of the sex bias (log_2_FC) and the sequence divergence rate (dN/dS) on sex-biased genes only, while controlling for the average expression level (CPM) of the gene, and GC-content (ppcor v.01 (Kim 2015), in R) in order to determine if genes with higher log_2_FC values have faster sequence divergence rates. Per gene GC content was calculated for the coding sequences from the *T. californicum* genome.

### Functional analysis of sex-biased genes

We performed Gene Ontology (GO) enrichment analysis using TopGO v.2.26.0 on sex-biased genes obtained with EdgeR (see above) at each developmental stage. We used a functional annotation derived from blasting sequences to *Drosophila melanogaster* database. Only GO terms with minimum ten annotated genes were used. Enrichment of terms was determined using a weighted Kolmogorov-Smirnov-like statistical test, equivalent to the gene set enrichment analysis (GSEA method). We applied the “elim” algorithm, which considers the Gene Ontology hierarchy, i.e. it first assesses the most specific GO terms, and then more general ones (Alexa *et al*. 2006). Our analysis focused on gene sets in the Biological Processes (BP) GO category. We considered terms as significant when p< 0.05. Enriched GO terms were then semantically clustered using ReviGO (Supek *et al*. 2011) to aid interpretation.

## Results

### Sex-biased gene expression increases during development in *T. californicum*

Sex-biased gene expression gradually increased during the three developmental stages, with 0.2% (26) of the expressed genes sex-biased at the hatchling stage, 4.7% (568) at the juvenile, and 20.3% (2485) at the adult stage (Fig. 2). There are significantly fewer sex-biased genes both at the hatchling stage compared to the juvenile stage (*χ*^2^_(1)_ = 498.11, p_adj._= 2.2 × 10^−16^) and at the juvenile compared to the adult stage (*χ*^2^_(1)_ = 1347, p_adj._= 2.2 × 10^−16^). In addition, the two effect sizes are similar; h_(hatchling-juvenile)_ = 0.34, h_(juvenile-adult)_ = 0.49 (Cohen 1988), supporting a gradual rather than abrupt increase. The hatchling stage had more female than male biased genes (*χ*^2^_(1)_=11.13, p= 0.0004), while the juvenile and adult stages had more male biased genes (*χ*^2^_juvenile (1)_ =467.1, *χ*^2^_adult (1)_ =85.915, p< 2.2 × 10^−16^) (Supplemental Table 4). All except one of the sex-biased genes at the two preadult stages had strong sex-bias (> 2 FC), while at the adult stage 77% had strong sex-bias (> 2 FC) (Fig. 2, see also Supplemental Table 4 and Fig 8.). The intensity of sex-bias varied greatly during development, with 1137 genes showing a significant sex by development interaction.

**Fig. 2.**
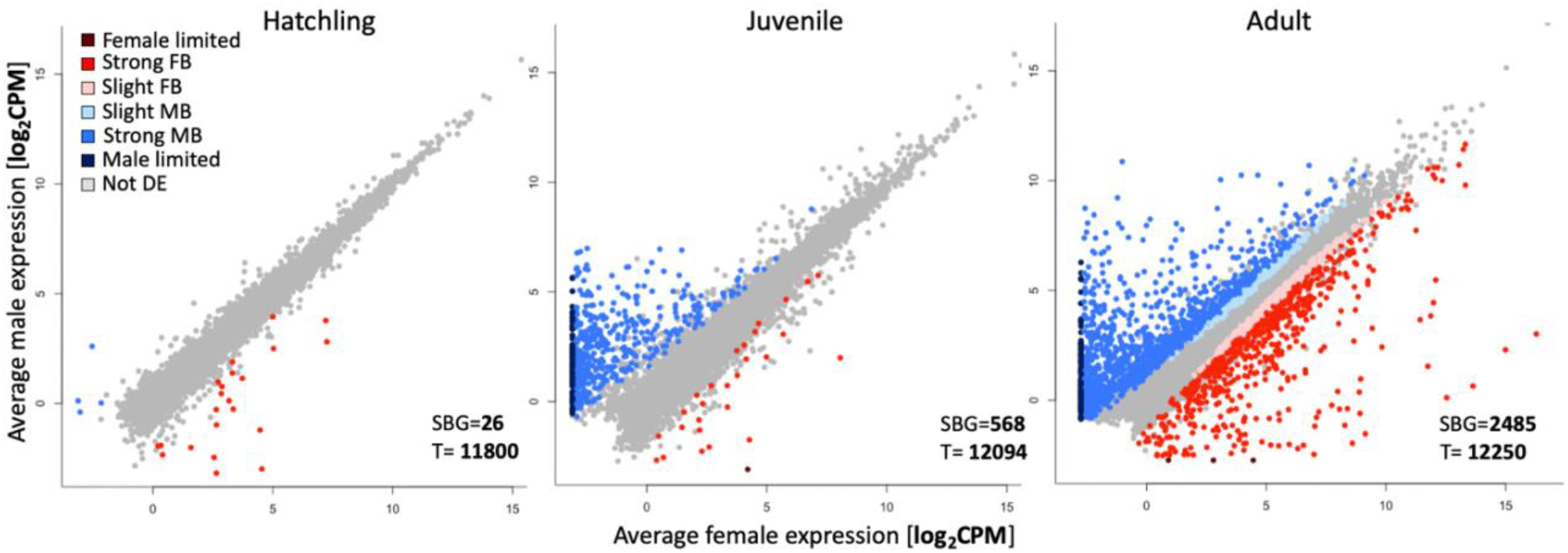
Gene expression (log_2_CPM) in *T. californicum* males and females at the hatchling (left), juvenile (middle) and adult stage (right). The number of differentially expressed genes (SBG) is shown at the bottom right corner of each plot, as well as the total number of expressed genes at each stage (T). Genes are classified based on their sex-bias into seven categories: “slight FB”- female bias (< 2 FC), “strong FB”- female bias (> 2 FC), “female limited”- with no expression in males, “slight MB”- male bias (< 2 FC), “strong MB”- male bias (> 2 FC), “male limited”- no expression in females, “Not DE”- not differentially expressed genes.

To characterize the dynamics of sex-biased expression during development, we then classified the 2671 genes which are sex-biased at one or more developmental stages into 13 categories, dependent on their expression patterns during development (Supplemental Fig. 1). This classification showed that sex-biased genes are added gradually during development, with genes sex-biased at earlier stages generally remaining sex-biased in the same direction at later stages (Supplemental Fig. 1 and 2). Only a single gene shifted from male-biased to female-biased expression (Supplemental Fig. 1). Out of 26 sex-biased genes at the hatchling stage, 18 (69%) were also significantly sex-biased in at least one of the later stages, with 9 (35%) remaining sex-biased throughout development (Fig. 3), while out of 568 sex-biased genes at the juvenile stage, 390 (69%) stayed sex-biased at the adult stage (Fig. 3). This classification is conservative, as it depends on the sex-biased gene expression detection threshold, meaning sex-biased gene expression may be missed in some stages, inflating the number of differences we see. This is supported by the fact that only around 65% of genes that are sex-biased in two stages show a significant sex by development stage interaction (Supplemental Table 8, also see Supplemental Fig. 1). Furthermore, the fold changes of sex-biased genes were strongly correlated between all developmental stages (ρ = 0.75 - 0.9 Supplemental Fig. 3).

**Fig. 3.**
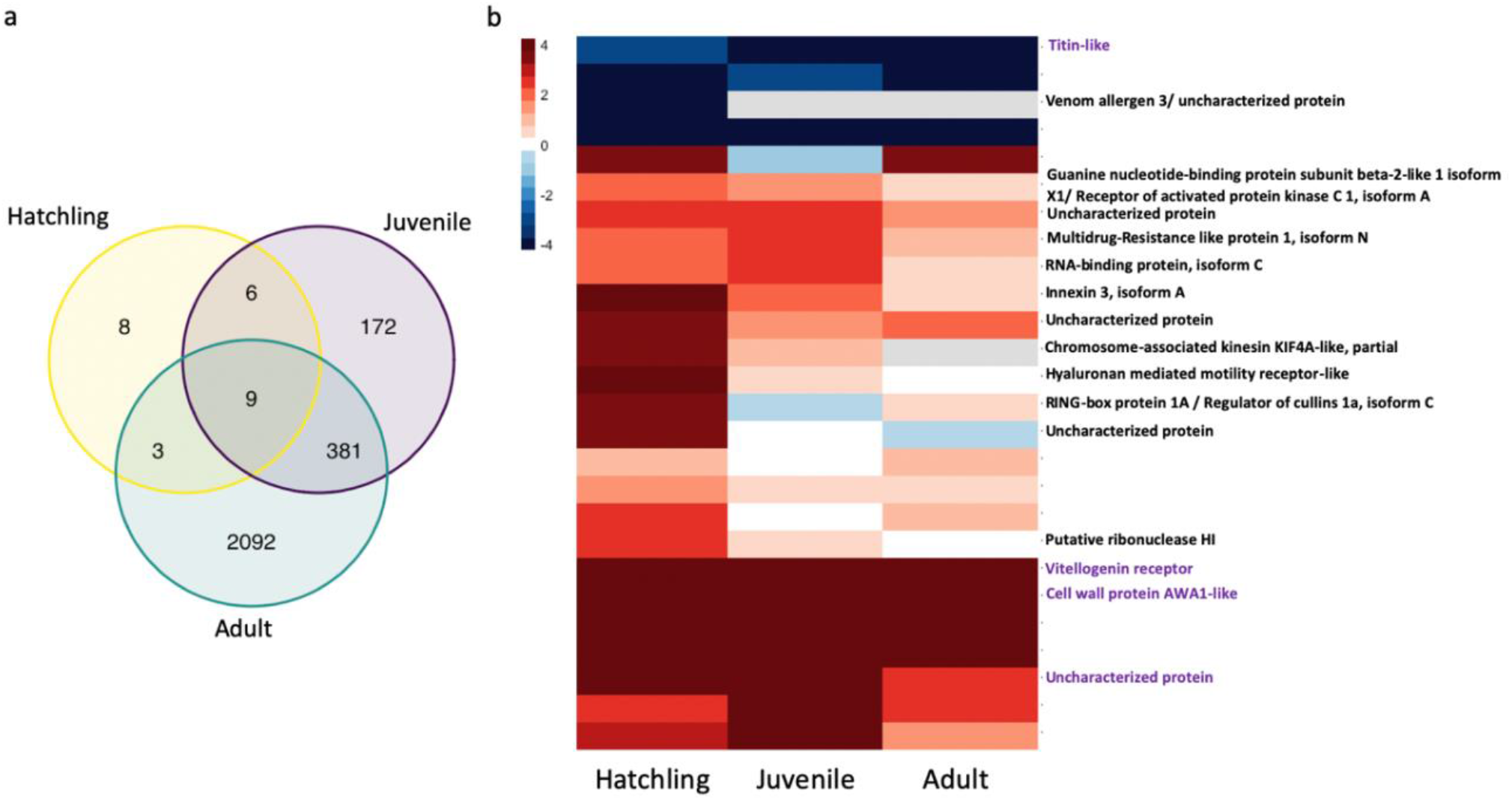
Sex-biased genes shared across development. (a) Venn-diagram of sex-biased genes in three developmental stages: hatchling (yellow), juvenile (purple), and adult (green). The number of genes shared between all three stages was greater than expected by chance (observed overlap= 9, expected overlap= 0.21, Exact test of multi set interactions: P_adj._= 5 × 10^−13^). Note that all pairwise overlaps also contained more genes than expected by chance (detailed results in Supplemental Table 7). (b) A heatmap showing the 26 sex-biased genes at the hatchling stage with their annotations and their expression in three developmental stages. Genes in red are female-biased, blue are male-biased. Saturation of the colors increases with the log_2_FC. Genes without a label have no annotation, annotation in purple is for genes that were sex-biased in all three stages.

Sex-biased genes at the hatchling stage are particularly interesting because they reveal sexual differentiation prior to visible morphological differences. We therefore looked for functional annotations of the 26 genes sex-biased at the hatchling stage in the *T. californicum* reference genome (Jaron *et al*. 2020). However, only 16 out of the 26 sex-biased genes had functional annotations and only one (vitellogenin receptor) had a clear link to sexual differentiation (Fig. 3b). While key genes known to play a role in insect sex-determination and differentiation pathways (*doublesex, transformer-2* and *sex-lethal*) were expressed at the hatchling stage, *doublesex* had very low expression, preventing analysis of sex-bias, and *transformer-2* and *sex-lethal* did not feature sex-biased expression.

### Sex-biased genes are enriched for development-related processes

Genes sex-biased at the hatchling stage were enriched for GO-terms related to developmental processes (e.g., “regulation of cell development”, “regulation of cell proliferation”, “cuticle development”), and GO-terms related to female specific functions such as oogenesis (“oocyte construction”, “oocyte development”) (Supplemental Fig. 5 a, and Supplemental Table 10). At the juvenile stage, sex-biased genes were enriched for GO-terms related to metabolic processes, in particular to catabolism (e.g., “lipid catabolic processes”, “cellular catabolic processes”, “regulation of catabolic processes”) (Supplemental Fig. 5 b, and Supplemental Table 11). At the adult stage, sex-biased genes were enriched for GO-terms related to diverse metabolic and physiological processes with no clear association to sexual differentiation (Supplemental Fig. 5 c, and Supplemental Table 12); however, several terms were related to chemosensory and olfactory behavior, which may play a role in mate detection. Furthermore, sex-biased genes at the adult stage were enriched for pigmentation (e.g., “developmental pigmentation”, “eye pigmentation”), which is a sexually dimorphic trait in *T. californicum* (Sandoval 2008).

### Sex-biased genes have more stage-specific expression

Expression levels of sex-biased genes tend to be specific to only one developmental stage while unbiased genes have a more constant expression level across development (Fig. 4, Supplemental Fig. 4). Male-biased genes have the most stage specific expression in both males and females. Female-biased genes are only slightly more stage specific than un-biased genes, but the trend is consistent and the difference is significant notably in both male and female adults.

**Fig. 4.**
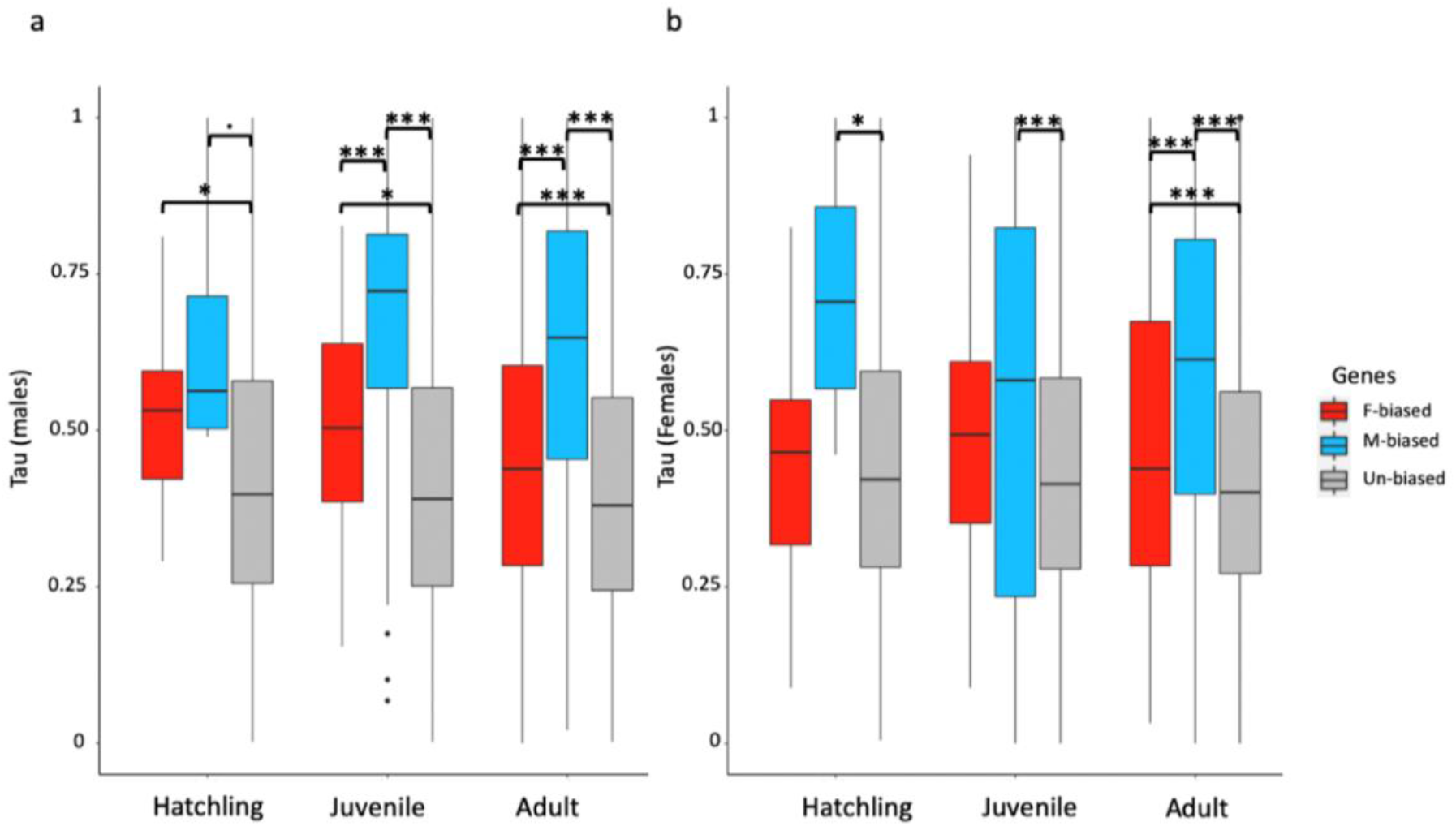
Tau index of gene expression at three developmental stages in males (a) and females (b). Tau ranges from zero (similarly expressed during development) to one (gene expressed in only one stage). Three gene categories are depicted with different colors; female-biased in red, male-biased in blue, and un-biased in grey. Boxplots represent the median, lower and upper quartiles, and whiskers the minimum and maximum values (in the limit of 1.5x interquartile range). Adjusted p values of Wilcoxon rank sum tests are summarized above the box plots (*** = p < 0.001, * = p < 0.05, · p=0.06).

### Sex-biased genes have faster sequence evolution rates

Male-biased genes have faster rates of sequence evolution (dN/dS) compared to both un-biased and female-biased genes, at juvenile and adult stages (Fig. 5). Female-biased genes do not evolve significantly faster than un-biased genes, at any of the three developmental stages.

**Fig. 5.**
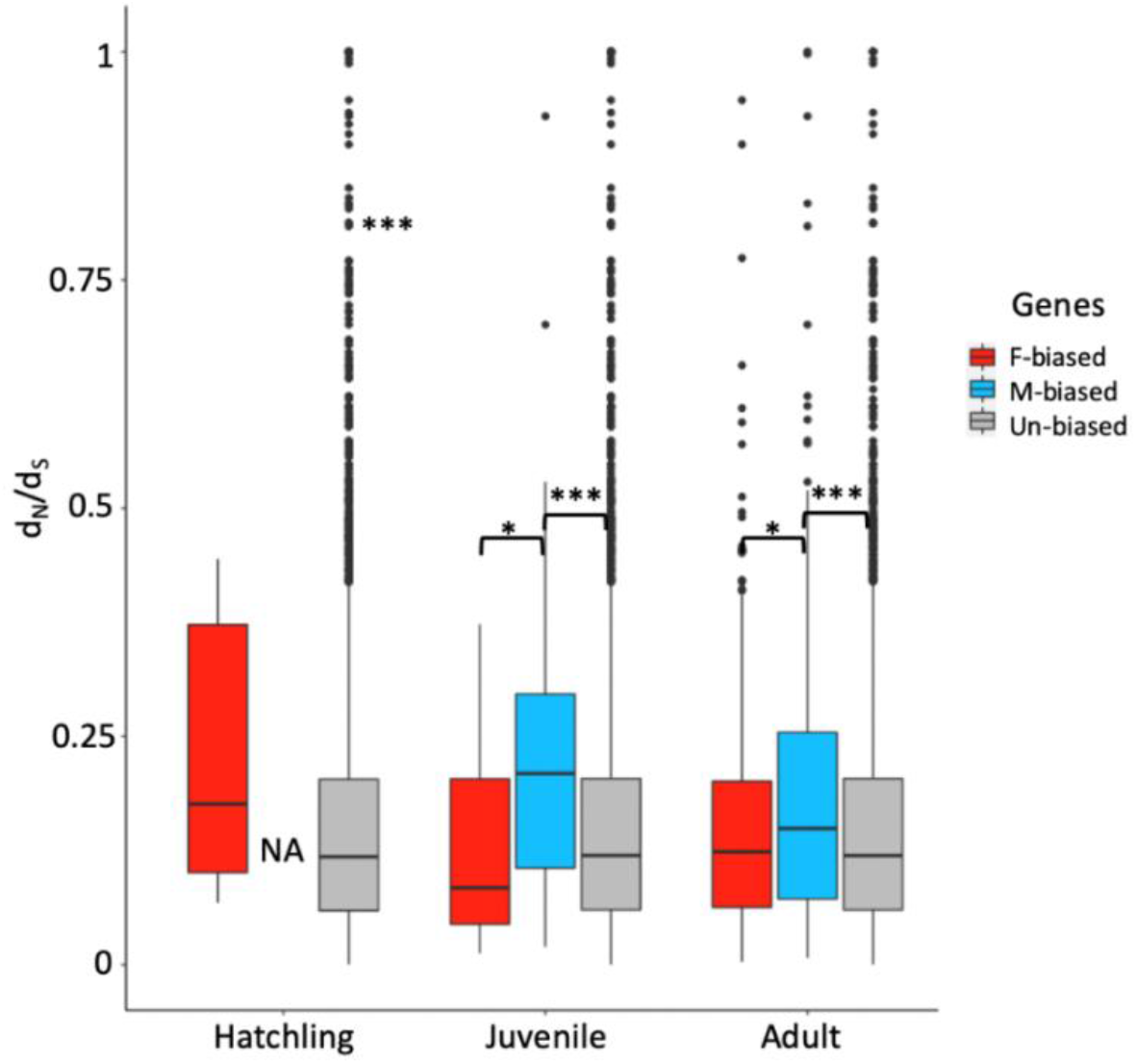
Divergence rates (the ratio of nonsynonymous to synonymous substitutions, dN/dS), at three developmental stages; hatchling, juvenile and adult stage. Boxplots represent the median, lower and upper quartiles, and whiskers the minimum and maximum values (in the limit of 1.5x interquartile range). Adjusted p values of Wilcoxon rank sum tests are summarized above the box plots (*** = p < 0.001, * = p < 0.05). Note that the four male-biased genes at the hatchling stage had no identified ortholog in other *Timema* species and therefore no associated dN/dS value.

We performed partial correlation analyses to test the effect of sex bias strength on sequence evolutionary rate, controlling for the average expression level of the gene, and GC-content (Supplemental Table 6). For both male-biased and female-biased genes, stronger sex-bias is associated with faster sequence evolution at the adult stage (Fig. 6). Partial correlations are not significant at juvenile or hatchling stages.

**Fig. 6.**
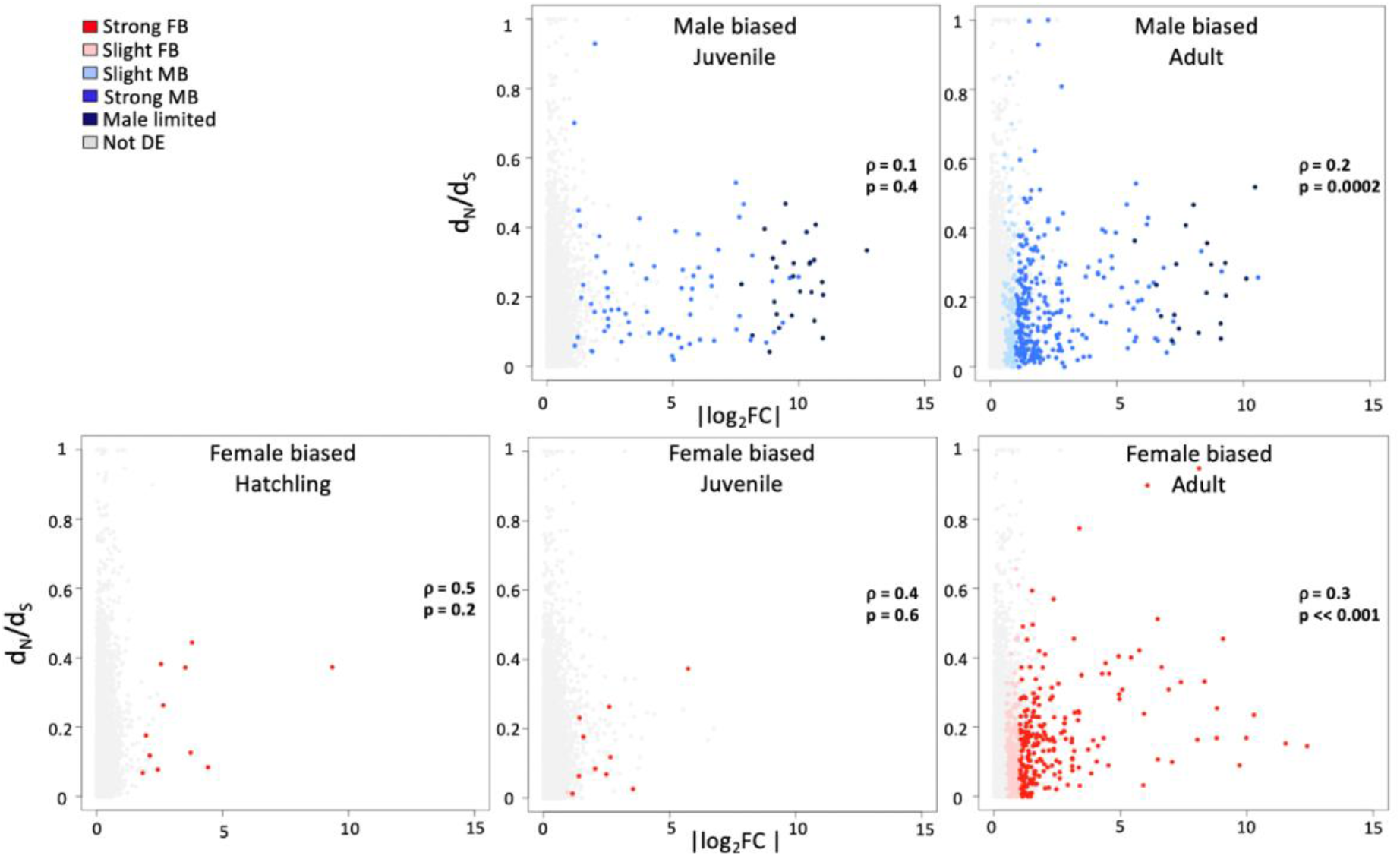
Partial Spearman’s rank correlations between the divergence rate of genes, dN/dS, and their strength of the sex bias |log_2_FC| at three developmental stages. Note that all genes are displayed on each plot but only male-respectively female-biased ones are used for correlation tests. Adjusted p values and partial correlation coefficients are shown in each plot.

### Sex-biased gene expression increases abruptly in adult *D. melanogaster*

In order to compare the dynamics of sex-biased gene expression during development between hemimetabolous and holometabolous insects, we reanalyzed publicly available data from *D. melanogaster* (Ingleby *et al*. 2016), using the same pipeline as for *T. californicum*. This experiment was chosen for comparison as it has a similar design, i.e. whole-body RNA-seq data from several developmental stages. In addition, although *D. melanogaster* from this experiment were lab-reared, they had a similar between sample variance as *T. californicum* (biological coefficient of variation at the adult stage: *D. melanogaster* = 0.274, *T. californicum* = 0.348).

In contrast to the gradual increase in the amount of sex-biased expression during development observed in *T. californicum*, sex-biased gene expression increased abruptly at the adult stage in *D. melanogaster* (Fig. 7 and Fig. 8.), from 22% to 84% of sex-biased genes (20% to 69% restricting to > 2 FC) (Fig. 8). There were significantly fewer sex-biased genes both at the larval compared to the pupal stage (*χ*^2^_(1)_ = 19.445, p_adj._ = 5.18 × 10^−6^) and at the pupal compared to the adult stage (*χ*^2^_(1)_ = 9607.2, p _adj._= 4.40 × 10^−16^). The two effect sizes were very different, with a more extensive shift from pupa to adult (h _larva-pupa_= 0.06, h _pupa-adult_ = 1.35) supporting the abrupt increase. Moreover, adult *D. melanogaster* had a significantly larger proportion of sex-biased genes than *T. californicum* (*χ*^2^_(1)_ =10061, p< 2.2 × 10^−16^). Overall, at each developmental stage, there were more male than female-biased genes (Supplemental Table 5). The majority of sex-biased genes had a strong sex bias at every developmental stage (> 2 FC: 83%, 94% and 82%, in larval, pupal, and adult stage, respectively) (Supplemental Table 5). Out of 2113 sex-biased genes at the larval stage, 1262 (60%) were significantly sex-biased in at least one of the later stages, with 1226 (58%) remaining sex-biased throughout development (Fig. 7b), while out of 2643 sex-biased genes at the pupal stage, 2423 (92%) stayed sex-biased at the adult stage (Fig. 7b). The fold changes of sex-biased genes were also low to moderately correlated between larvae and pupae or adults (ρ = 0.30 and 0.69; Supplemental Fig. 3) but strongly correlated between pupae and adults (ρ = 0.85; Supplemental Fig. 3).

**Fig. 7.**
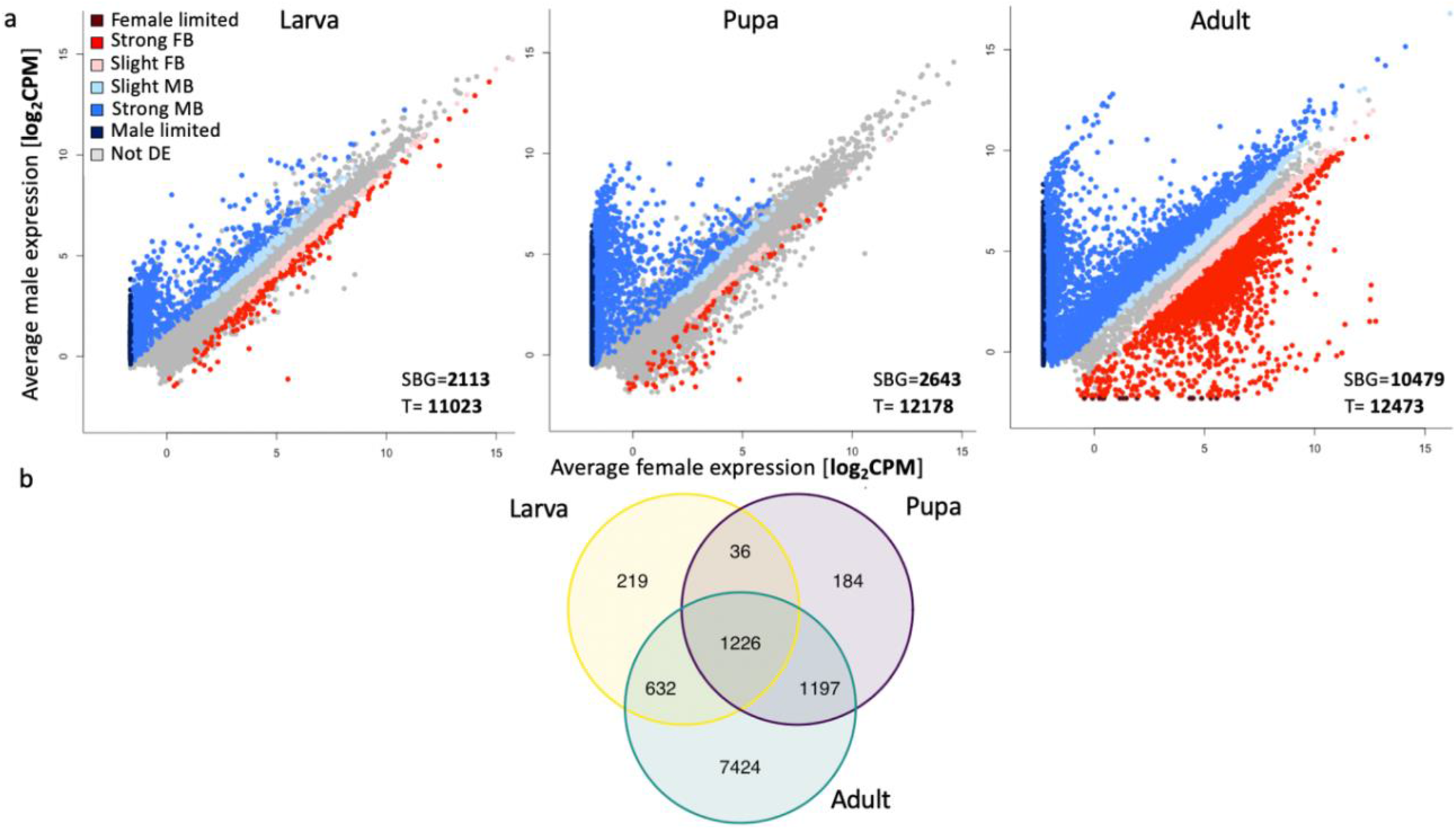
**(a)** Gene expression (log_2_CPM) in males and females at the larval (left), pupal (middle) and adult stage (right). The number of differentially expressed genes (SBG) is shown at bottom right corner of each plot, as well as the total number of expressed genes at each stage (T). Genes are classified based on their sex-bias into seven categories: “slight FB”- female bias (< 2 FC), “strong FB”- female bias (> 2 FC), “female limited”- with no expression in males, “slight MB”- male bias (< 2 FC), “strong MB”- male bias (> 2 FC), “male limited”- no expression in females, “Not DE”- not differentially expressed genes. **(b)** Venn-diagram of sex-biased genes in three developmental stages: larval (yellow), pupal (purple), and adult (green). The number of genes shared between all three stages was greater than expected by chance (observed overlap = 1226, expected overlap = 334.7), Exact test of multi set intersections: p-value ≈ 0).

**Fig. 8.**
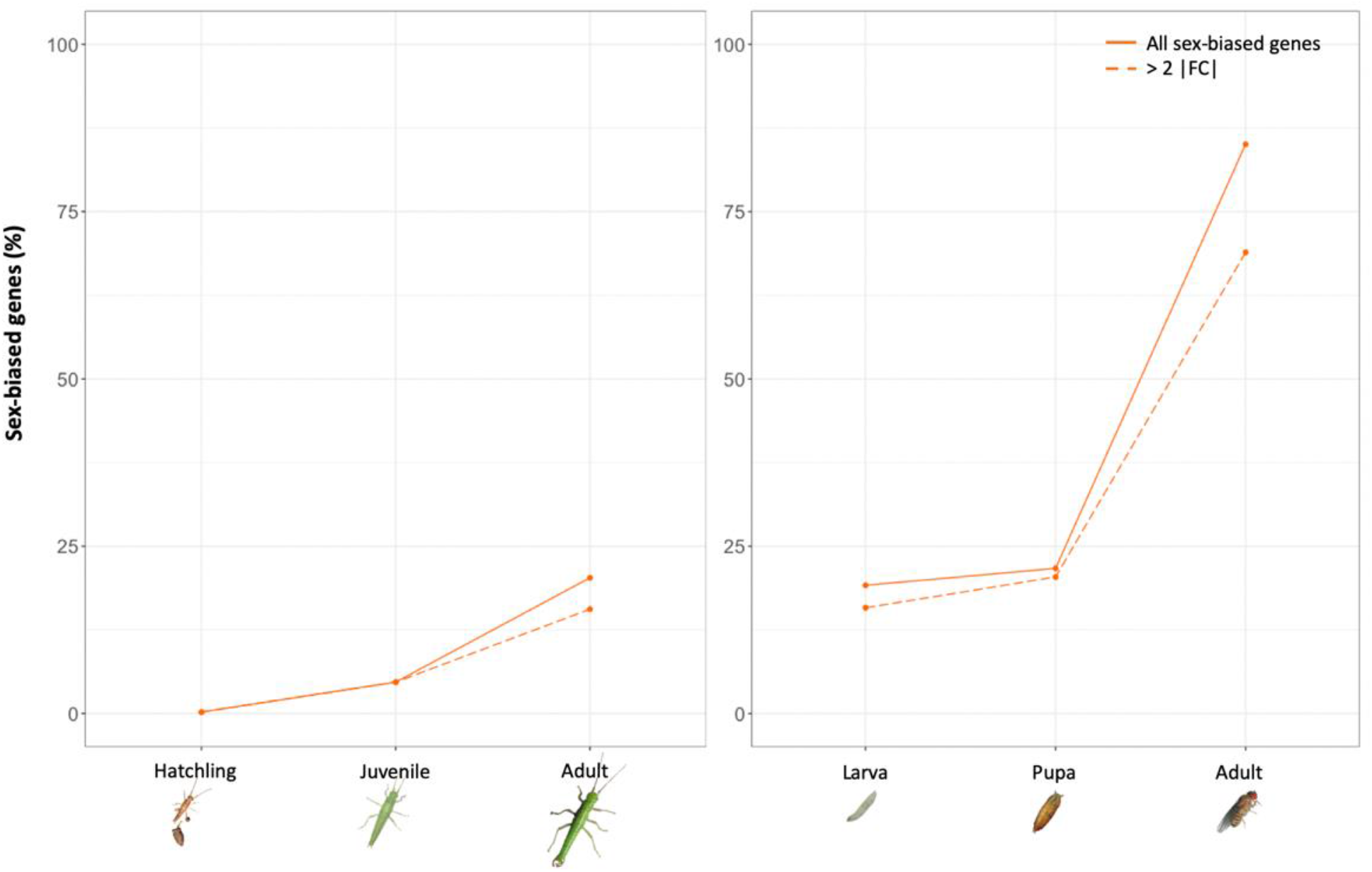
Percentage of sex-biased genes (SBG) in *T. californicum* (left panel) and *D. melanogaster* (right panel) at three developmental stages. Note that the immature developmental stages in the two species are not homologous but serve to illustrate the gradual versus abrupt increase in sex-biased gene expression in adults.

## Discussion

In the stick insect *T. californicum*, gene expression differs between males and females before morphological sexual dimorphism. During development, most of the sex-biased genes are added gradually and cumulatively, meaning that once a gene gains sex-biased expression in *T. californicum*, it generally remains sex-biased at later developmental stages. In addition, genes were consistent in their sex-bias direction; only a single gene shifted from male- to female-biased expression. Surprisingly, such a gradual and consistent addition of sex-bias does not appear to be a general pattern over animal development. For example, in humans and mosquitos, some stages have bursts of sex-biased gene expression, suggesting that specific developmental stages contribute discretely to sexual differentiation, and that sexual phenotypes are not gradually established (Magnusson *et al*. 2011; Shi *et al*. 2016).

Although only 26 genes were sex-biased in *T. californicum* at the hatchling stage, they all had a strong sex-bias and nine of them stayed significantly sex-biased at both the juvenile and adult stages. As such these 26 genes likely represent key genes involved in the sexual differentiation of *Timema*. Most of the sex-biased genes at the hatchling stage were female-biased. This might suggest that the process of building a female phenotype begins earlier in development than the one of building a male phenotype. Later developmental stages had more male-biased than female-biased genes. A similar shift from female- to male-bias in gene expression during development was found in *Nasonia* jewel wasps, where little male-biased expression was detected until the pupal stage, during the activation of spermatogenesis. Similar to *Timema*, the shift was interpreted as a different developmental timing of the two sexes (Rago *et al*. 2020). In *Timema*, male-biased genes were also expressed in a relatively stage-specific manner, in contrast to a more constant expression of female-biased genes. This further supports the idea that *Timema* individuals first develop along the female trajectory, i.e. that the female phenotype is the “default” in *Timema*, and may indicate greater pleiotropy of female-biased genes. Additional support for this interpretation comes from the lower sequence divergence rates of female-biased than male-biased genes, implicating stronger selective constraints on female- than male-biased genes. In addition, we observed that genes with greater sex-bias have faster sequence divergence rates. This is largely due to relaxed purifying selection, with positive selection contributing little to the accelerated evolutionary rate of sex-biased genes. These findings are consistent with the idea that sex-biased genes evolve from genes with few evolutionary constraints and that are relatively “dispensable” (Mank and Ellegren 2009; Catalan *et al*. 2018).

Sex-biased genes in *Timema* hatchlings were enriched for processes related to oogenesis. Moreover, one of the most consistently and strongly female-biased genes across development was annotated as vitellogenin receptor. In insects, vitellogenin receptor transfers yolk protein precursors into the oocytes and is necessary for oocyte and early embryo development (Raikhel and Dhadialla 1992; Cho and Raikhel 2001). The role of these processes in *Timema* hatchlings remain to be investigated, but vitellogenin receptor expression may indicate a very early onset of oocyte development in *Timema*, consistent with studies in other stick insects (Taddei *et al*. 1992). Sex-biased genes in juvenile and adult stages were not enriched for processes obviously related to sexual traits. However, some processes such as pigmentation and chemosensory/olfactory behavior could have a sex-related role. Pigmentation is a sexually dimorphic trait in *Timema* (Sandoval 2008), while chemosensory and olfactory behaviors are important for mate recognition and mating (Nosil *et al*. 2007; Schwander *et al*. 2013). We also checked the expression of two key genes involved in insect sex-determination pathways: *transformer* and *doublesex*. In holometabolous insects, *transformer* acts as a splicing regulator of *doublesex*, a sex differentiation master switch gene (Geuverink and Beukeboom 2014). The *doublesex-transformer* role in sex-differentiation is conserved among Diptera, Coleoptera and Hymenoptera (Verhulst *et al*. 2010; Wexler *et al*. 2019), and likely play the same role in sex differentiation in hemimetabolous insects (Zhuo *et al*. 2018; Wexler *et al*. 2019). In *T. californicum*, we found that *doublesex* was expressed in all three developmental stages, but its expression was too low to examine sex-bias. *Transformer* is not annotated in the available *T. californicum* genome; a *transformer-2* homolog is annotated but did not feature sex-bias at any of the three developmental stages. As such, whether or not these classic sex-determining genes influence sexual differentiation in *Timema* is unclear, and requires future studies.

We hypothesized that the dynamics of sex biased gene expression would be strongly affected by hemimetabolous vs holometabolous development in insects, with a gradual increase of sex-biased gene expression during development in the former and an abrupt increase at the adult stage in the latter. Hemimetabolous insects such as *T. californicum* have multiple nymphal stages that progressively resemble the adult stage, together with a gradual increase in sexual dimorphism. On the other hand, a holometabolous insect like *D. melanogaster* has more monomorphic larval stages and pupae, with no resemblance to the sexually dimorphic adult stage. The change in sexually dimorphic gene expression across development indeed mirrored these different developmental patterns, with the proportion of sex-biased genes increasing gradually in *T. californicum* as sexual dimorphism became more pronounced, whereas sexually dimorphic gene expression increased abruptly at the adult stage in *D. melanogaster*.

Although the change in sexually dimorphic gene expression across development fits with our expectations, we were surprised to find a much higher proportion of sex-biased genes in *D. melanogaster* than in *T. californicum* overall. This difference is clearest at the adult stage when most of the expressed genes in *D. melanogaster* were sex-biased (84%), compared to only 20% in *T. californicum*. Similar to *Drosophila*, other holometabolous insects also featured sex-bias for more than half of the genes expressed at the adult stage (Baker *et al*. 2011; Rago *et al*. 2020), while all studied hemimetabolous insects had a much smaller proportion of sex-biased genes (< 10%) (Pal and Vicoso 2015). Why there is such a difference in the prevalence of sex-biased genes between species with different developmental strategies is still an open question. One potential explanation is that the proportion of sex-biased genes at the adult stage may be less constrained under holo- than hemimetabolous development. For example, the complete reorganization of cells and tissues during metamorphosis in holometabolous insects may allow for more drastic phenotypic changes between the sexes, while incomplete metamorphosis constrains the evolution of distinct sexual phenotypes. Data on additional species and specific tissues rather than whole bodies are required to confirm that this difference is a general phenomenon and not simply due idiosyncrasies of the few species yet investigated. Holometabolous insects studied are often established lab-models, with high fecundity, and may thus be characterized by comparatively large gonads, which would result in large proportions of sex-biased genes in adults. By contrast, all hemimetabolous insects studied for sex-biased expression are non-model species. However, the choice of model species is unlikely to be the sole explanation for the different amount of sex bias in holo- and hemimetabolous species. Indeed, in the cricket *Gryllus bimaculatus* and in *Timema* stick insects, even the most sexually differentiated tissue (gonads) feature sex-bias of fewer than 30% of genes (Parker *et al*. 2019; Whittle *et al*. 2020), suggesting that extensive differences are not completely driven by large gonads. Finally, while whole body analysis such as we have performed here provides a window into ontogeny, it will necessarily miss tissue specific regulation of sex-biased gene expression (Montgomery and Mank 2016). Future studies should therefore study sex-biased gene expression during development by examining a panel of different tissues or cell types.

Overall, our results describe the dynamics of sex-biased gene expression during development in a hemimetabolous insect. Generating sexually dimorphic phenotypes is a developmental process, and we show that dynamics of sex-biased gene expression mirror this development with sex-biased genes being added gradually during development, and with the majority of genes sex-biased at early stages remaining sex-biased in later stages.

## Data accessibility

Data are available online: *Timema californicum* raw reads have been deposited in SRA under PRJNA678950 bioproject, with accession codes SRR13084978-SRR13085001 (Supplemental Table 9). *Timema californicum* mapped read counts are provided in Supplemental Table 13 and fold changes in Supplemental Table 14. *Drosophila melanogaster* mapped read counts are provided in Supplemental Table 15.

## Supplementary material

Script and code are available online: DOI 10.5281/zenodo.5639300

## Acknowledgements

This study was supported by Swiss National Science Foundation Grant 31003A_182495. Version 6 and 7 of this preprint have been peer-reviewed and recommended by Peer Community In Evolutionary Biology (https://doi.org/10.24072/pci.evolbiol.100135).

## Conflict of interest disclosure

The authors of this preprint declare that they have no financial conflict of interest with the content of this article. Tanja Schwander and Marc Robinson-Rechavi are recommenders for PCI.

## Appendix

Link to the supplementary material: https://www.biorxiv.org/content/10.1101/2021.01.23.427895v7.supplementary-material

Appendix 1- Supplemental Figs 1-5, Supplemental Tables 1-9

Appendix 2- GO-terms hatchling stage- Supplemental table 10

Appendix 3- GO-terms juvenile stage- Supplemental table 11

Appendix 4- GO-terms adult stage- Supplemental table 12

Appendix 5- *T. californicum* mapped read counts- Supplemental Table 13

Appendix 6- *T. californicum* fold-changes- Supplemental Table 14

Appendix 7- *D. melanogaster* mapped read counts- Supplemental Table 15

## Notes

### Competing Interest Statement

The authors have declared no competing interest.

### Summary of Updates

Version 6 and 7 of this preprint has been peer-reviewed and recommended by Peer Community In Evolutionary Biology (https://doi.org/10.24072/pci.evolbiol.100135).

