## Supplemental tables and figures for "Dynamics of sex-biased gene expression during development in the stick insect *Timema californicum*"

### Supplemental material

**Supplemental table 1|** The sex of *Timema* hatchlings was determined via X-linked microsatellite genotyping using the indicated primer sequences **(a)** and PCR conditions **(b)**

**a)**

| Usat ID | Primer ID | Forward 5'-3' | Primer ID | Reverse 5'-3' |
|---|---|---|---|---|
| **19** | Cm_19F | CATCGAGTGTTGCTAAGAATGG | Cm_19R | AGACTTTAACCCAGGCCCAG |
| **20** | Cm_20F | TGTGCGAGCAAATGAGGTTG | Cm_20R | CGATGGCATGTCTTAAGCCC |
| **21** | Cm_21F | GCCAATCGAAGCGGGTATTG | Cm_21R | GTATGGAGCGAGTGACAAAGC |
| **24** | Cm_24F | TCGCTGCACATTTCTACAAAC | Cm_24R | CTGAATACCGGCAGTGAAGG |

**b)**

| Cycling conditions | | | |
|---|---|---|---|
| | **Temp** | **Time** | **Cycles** |
| **Denaturation initial** | 95°C | 15 min | |
| **Denaturation** | 94°C | 30sec | |
| **Annealing** | 57°C | 90sec | 35x |
| **Extension** | 72°C | 1 min | |
| **Final extension** | 60°C | 30min | |
| | 4°C | 10 min | |
| | 12°C | xxx | |

**Supplemental table 2|** Genotypes for four X-linked microsatellite loci (Marker: 19, 20, 21 and 24) recorded in 15 male and 15 female adults and nine hatchlings. Heterozygotes are highlighted in grey.

| Stage | Sample ID | Sex | Marker 19 | | Marker 20 | | Marker 21 | | Marker 24 | |
|---|---|---|---|---|---|---|---|---|---|---|
| **Adult** | HM217 | F | 376 | 386 | 390 | 390 | 281 | 312 | 197 | 219 |
| | HM218 | F | 351 | 383 | 390 | 390 | 277 | 312 | 200 | 200 |
| | HM219 | F | 353 | 370 | 394 | 394 | 281 | 308 | 197 | 203 |
| | HM220 | F | 353 | 390 | 390 | 394 | 277 | 308 | 191 | 191 |
| | HM221 | F | 374 | 403 | 394 | 400 | 304 | 312 | 191 | 197 |
| | HM222 | F | 388 | 395 | 396 | 412 | 277 | 308 | 197 | 206 |
| | ReSeq_Cm02 | F | 351 | 351 | 390 | 398 | 288 | 308 | 203 | 203 |

| | | | | | | | | | | |
|---|---|---|---|---|---|---|---|---|---|---|
| | ReSeq_Cm04 | F | 353 | 374 | 392 | 394 | 292 | 312 | 191 | 197 |
| | ReSeq_Cm06 | F | 374 | 393 | 390 | 394 | 277 | 281 | 197 | 197 |
| | ReSeq_Cm08 | F | 353 | 392 | 396 | 396 | 296 | 304 | 197 | 197 |
| | ReSeq_Cm10 | F | 351 | 360 | 392 | 394 | 308 | 312 | 203 | 203 |
| | ReSeq_Cm12 | F | 351 | 393 | 394 | 394 | 277 | 312 | 197 | 206 |
| | ReSeq_Cm14 | F | 355 | 355 | 394 | 394 | 288 | 308 | 191 | 203 |
| | ReSeq_Cm16 | F | 370 | 378 | 394 | 396 | 304 | 304 | 197 | 200 |
| | ReSeq_Cm18 | F | 378 | 390 | 394 | 396 | 277 | 292 | 197 | 197 |
| | HM157 | M | 351 | | 392 | | 304 | | 197 | |
| | HM157b | M | 369 | | 390 | | 274 | | 197 | |
| | HM158 | M | 380 | | 394 | | 281 | | 197 | |
| | HM159 | M | 348 | | 392 | | 324 | | 194 | |
| | HM160 | M | 386 | | 394 | | 292 | | 206 | |
| | HM161 | M | 376 | | 394 | | 292 | | 203 | |
| | HM162 | M | 393 | | 396 | | 308 | | NA | |
| | HM163 | M | 351 | | 390 | | 304 | | NA | |
| | HM164 | M | 351 | | 394 | | 308 | | 197 | |
| | HM165 | M | 382 | | 394 | | 304 | | 201 | |
| | HM166 | M | 376 | | 392 | | 292 | | 200 | |
| | Tcm_M_Hm01 | M | 374 | | 396 | | 308 | | 206 | |
| | Tcm_M_Hm02 | M | 390 | | 410 | | 324 | | 197 | |
| | Tcm_M_Hm03 | M | 351 | | 408 | | 296 | | 195 | |
| | Tcm_M_Hm04 | M | NA | | NA | | 304 | | NA | |
| **Hatchling** | Tim_Ha09 | F | 353 | 356 | 399 | | 296 | | 196 | 199 |
| | Tim_Ha10 | M | 353 | | 405 | | NA | | 196 | |
| | Tim_Ha11 | M | 353 | | 399 | | NA | | 199 | |
| | Tim_Ha14 | M | 355 | | 397 | | NA | | 199 | |
| | Tim_Ha15 | F | 351 | 355 | 388 | 397 | NA | | 196 | |
| | Tim_Ha16 | M | 353 | | 399 | | NA | | 199 | |
| | TIM_HA12 | F | 353 | 356 | 399 | 405 | 295 | | NA | |
| | TIM_HA13 | M | 353 | | 388 | | 299 | | NA | |
| | TIM_HA18 | F | 353 | 356 | NA | NA | 295 | | NA | |

**Supplemental table 3|** – *D. melanogaster* samples, and their accession numbers

| **Accession** | **Sample ID** | **Sex** | **Stage** |
|---|---|---|---|
| SRR3092025 | FL.H169.1 | Female | Larva |
| SRR3091999 | ML.H8.1 | Male | Larva |
| SRR3092040 | FL.H60.3 | Female | Larva |

| | | | |
|---|---|---|---|
| SRR3092044 | FL.H74.3 | Female | Larva |
| SRR3091997 | ML.H75.2 | Male | Larva |
| SRR3091988 | ML.H55.3 | Male | Larva |
| SRR3092052 | FL.H94.2 | Female | Larva |
| SRR3091990 | ML.H60.2 | Male | Larva |
| SRR3092081 | FP.H8.2 | Female | Pupa |
| SRR3092085 | FP.H94.3 | Female | Pupa |
| SRR3092079 | FP.H75.3 | Female | Pupa |
| SRR3138724 | MP.H163.2 | Male | Pupa |
| SRR3138731 | MP.H184.3 | Male | Pupa |
| SRR3092055 | FP.H163.1 | Female | Pupa |
| SRR3138728 | MP.H169.3 | Male | Pupa |
| SRR3138734 | MP.H196.3 | Male | Pupa |
| SRR3092076 | FA.H184.3 | Female | Adult |
| SRR3092017 | FA.H8.3 | Female | Adult |
| SRR3091979 | MA.H94.1 | Male | Adult |
| SRR3091965 | MA.H55.3 | Male | Adult |
| SRR3092011 | FA.H74.3 | Female | Adult |
| SRR3091970 | MA.H74.2 | Male | Adult |
| SRR3091994 | FA.H55.2 | Female | Adult |
| SRR3091973 | MA.H75.1 | Male | Adult |

**Supplemental table 4|**- *Timema californicum*; Filtered genes (Number of genes filtered for low expression), kept (genes included for differential gene expression analysis), Fb (female-biased genes), Mb (male-biased genes), total SBG (sum of the male and female-biased genes), %SBG (percentage of sex-biased genes), % (>1 $log_2FC$) (percentage of genes with strong sex bias). Genes are classified based on their sex-bias into seven categories: “slight FB”- female bias (<1 $log_2FC$), “strong FB”- female bias (≥1 $log_2FC$), “female limited”- with no expression in males, “slight MB”- male bias (<1 $log_2FC$), “strong MB”- male bias (≥1 $log_2FC$), “male limited”- no expression in females, “Not DE”- not differentially expressed genes.

| | Filtered genes | Kept | Fb | Mb | total SBG | %SBG | % (>1 log2FC) | Slight FB | Slight MB | Strong FB | Strong MB | Female limited | Male limited | NotDE |
|---|---|---|---|---|---|---|---|---|---|---|---|---|---|---|
| **hatchling** | 2763 | 11800 | 4 | 22 | 26 | 0.22 | 0.22 | 0 | 0 | 22 | 4 | 0 | 0 | 11774 |
| **juvenile** | 2469 | 12094 | 29 | 539 | 568 | 4.70 | 4.69 | 1 | 0 | 27 | 417 | 1 | 122 | 11526 |
| **adult** | 2313 | 12250 | 1023 | 1462 | 2485 | 20.29 | 15.60 | 374 | 200 | 646 | 1139 | 3 | 123 | 9765 |

**Supplemental table 5|** - *Drosophila melanogaster*; Filtered genes (Number of genes filtered for low expression), kept (genes included for differential gene expression analysis), Fb (female-biased genes), Mb (male-biased genes), total SBG (sum of the male and female-biased genes), %SBG (percentage of sex-biased genes), % (>1 $log_2$FC) (percentage of genes with strong sex bias). Genes are classified based on their sex-bias into seven categories: "slight FB"- female bias (<1 $log_2$FC), "strong FB"- female bias (≥1 $log_2$FC), "female limited"- with no expression in males, "slight MB"- male bias (<1 $log_2$FC), "strong MB"- male bias (≥1 $log_2$FC), "male limited"- no expression in females, "Not DE"- not differentially expressed genes.

| | Filtered genes | Kept | Fb | Mb | total SBG | %SBG | %(>1 log2FC) | Slight FB | Slight MB | Strong FB | Strong MB | Female limited | Male limited | NotDE |
|---|---|---|---|---|---|---|---|---|---|---|---|---|---|---|
| **larva** | 6733 | 11023 | 399 | 1714 | 2113 | 19.17 | 15.81 | 202 | 168 | 197 | 1301 | 0 | 245 | 8910 |
| **pupa** | 5578 | 12178 | 151 | 2492 | 2643 | 21.70 | 20.41 | 63 | 94 | 88 | 1620 | 0 | 778 | 9357 |
| **adult** | 5283 | 12473 | 4497 | 5982 | 10479 | 84.00 | 68.91 | 1299 | 585 | 3181 | 4300 | 17 | 1097 | 1994 |

**Supplemental table 6|** Statistical results: partial correlations.

### Female-biased adult stage

**Estimate**

| n=443 gp=2 | dNdS | avr. exp. | GC | \|FC\| |
|---|---|---|---|---|
| dNdS | 1.00 | -0.03 | -0.26 | 0.32 |
| avr. exp | -0.03 | 1.00 | 0.03 | -0.19 |
| GC | -0.26 | 0.03 | 1.00 | 0.27 |
| \|FC\| | 0.32 | -0.19 | 0.27 | 1.00 |

**p value**

| | dNdS | avr. exp. | GC | \|FC\| |
|---|---|---|---|---|
| dNdS | 0.000 | 0.538 | 0.000 | 0.000 |
| avr. exp | 0.538 | 0.000 | 0.533 | 0.000 |
| GC | 0.000 | 0.533 | 0.000 | 0.000 |
| \|FC\| | 0.000 | 0.000 | 0.000 | 0.000 |

**statistic**

| | dNdS | avr. exp. | GC | \|FC\| |
|---|---|---|---|---|
| dNdS | 0.00 | -0.62 | -5.70 | 7.06 |
| avr. exp | -0.62 | 0.00 | 0.62 | -4.13 |
| GC | -5.70 | 0.62 | 0.00 | 5.95 |
| \|FC\| | 7.06 | -4.13 | 5.95 | 0.00 |

### Male-biased- adult stage

**Estimate**

| n=388 gp=2 | dNdS | avr. exp. | GC | \|FC\| |
|---|---|---|---|---|
| dNdS | 1.00 | 0.00 | -0.44 | 0.19 |
| avr. exp | 0.00 | 1.00 | 0.08 | -0.58 |
| GC | -0.44 | 0.08 | 1.00 | 0.22 |
| \|FC\| | 0.19 | -0.58 | 0.22 | 1.00 |

**p value**

| | dNdS | avr. exp. | GC | \|FC\| |
|---|---|---|---|---|
| dNdS | 0.000 | 0.975 | 0.000 | 0.000 |
| avr. exp | 0.975 | 0.000 | 0.099 | 0.000 |
| GC | 0.000 | 0.099 | 0.000 | 0.000 |
| \|FC\| | 0.000 | 0.000 | 0.000 | 0.000 |

**statistic**

| | dNdS | avr. exp. | GC | \|FC\| |
|---|---|---|---|---|
| dNdS | 0.00 | 0.03 | -9.70 | 3.81 |
| avr. exp | 0.03 | 0.00 | 1.65 | -14.06 |
| GC | -9.70 | 1.65 | 0.00 | 4.42 |
| \|FC\| | 3.81 | -14.06 | 4.42 | 0.00 |

### Female-biased juvenile

**Estimate**

| n=11 gp=2 | dNdS | avr. exp. | GC | \|FC\| |
|---|---|---|---|---|
| dNdS | 1.00 | -0.03 | -0.07 | 0.40 |
| avr. exp | -0.03 | 1.00 | 0.01 | -0.44 |
| GC | -0.07 | 0.01 | 1.00 | -0.40 |
| \|FC\| | 0.40 | -0.44 | -0.40 | 1.00 |

**p value**

| | dNdS | avr. exp. | GC | \|FC\| |
|---|---|---|---|---|
| dNdS | 0.000 | 0.932 | 0.932 | 0.573 |
| avr. exp | 0.932 | 0.000 | 0.974 | 0.240 |
| GC | 0.850 | 0.974 | 0.000 | 0.289 |
| \|FC\| | 0.286 | 0.240 | 0.289 | 0.000 |

**statistic**

| | dNdS | avr. exp. | GC | \|FC\| |
|---|---|---|---|---|
| dNdS | 0.00 | -0.09 | -0.20 | 1.15 |
| avr. exp | -0.09 | 0.00 | 0.03 | -1.28 |
| GC | -0.20 | 0.03 | 0.00 | -1.15 |
| \|FC\| | 1.15 | -1.28 | -1.15 | 0.00 |

### Male-biased juvenile

**Estimate**

| n= 94 gp=2 | dNdS | avr. exp. | GC | \|FC\| |
|---|---|---|---|---|
| dNdS | 1.00 | -0.10 | -0.34 | 0.08 |
| avr. exp | -0.10 | 1.00 | -0.08 | -0.38 |
| GC | -0.34 | -0.08 | 1.00 | -0.07 |
| \|FC\| | 0.08 | -0.38 | -0.07 | 1.00 |

**p value**

| | dNdS | avr. exp. | GC | \|FC\| |
|---|---|---|---|---|
| dNdS | 0.000 | 0.433 | 0.002 | 0.452 |
| avr. exp | 0.325 | 0.000 | 0.457 | 0.000 |
| GC | 0.001 | 0.457 | 0.000 | 0.491 |
| \|FC\| | 0.452 | 0.000 | 0.491 | 0.000 |

**statistic**

| | dNdS | avr. exp. | GC | \|FC\| |
|---|---|---|---|---|
| dNdS | 0.00 | -0.99 | -3.41 | 0.76 |
| avr. exp | -0.99 | 0.00 | -0.75 | -3.93 |
| GC | -3.41 | -0.75 | 0.00 | -0.69 |
| \|FC\| | 0.76 | -3.93 | -0.69 | 0.00 |

### Female-biased hatchling

**Estimate**

| n=14 gp=2 | dNdS | avr. exp. | GC | \|FC\| |
|---|---|---|---|---|
| dNdS | 1.00 | 0.50 | -0.52 | 0.51 |
| avr. exp | 0.50 | 1.00 | 0.11 | -0.05 |
| avr. exp. females | -0.52 | 0.11 | 1.00 | 0.33 |
| \|log2FC\| | 0.51 | -0.05 | 0.33 | 1.00 |

**p value**

| | dNdS | avr. exp. | GC | \|FC\| |
|---|---|---|---|---|
| dNdS | 0.000 | 0.174 | 0.151 | 0.162 |
| avr. exp | 0.174 | 0.000 | 0.777 | 0.905 |
| GC | 0.151 | 0.777 | 0.000 | 0.386 |
| \|FC\| | 0.162 | 0.905 | 0.386 | 0.000 |

**statistic**

| | dNdS | avr. exp. | GC | \|FC\| |
|---|---|---|---|---|
| dNdS | 0.00 | 1.51 | -1.61 | 1.56 |
| avr. exp | 1.51 | 0.00 | 0.29 | -0.12 |
| GC | -1.61 | 0.29 | 0.00 | 0.92 |
| \|FC\| | 1.56 | -0.12 | 0.92 | 0.00 |

**Supplemental table 7|** Exact test of multi set intersections of sex-biased genes between developmental stages in *T. californicum*

| Intersections | Degree | Observed Overlap | Expected Overlap | FE | P.value | P.adj |
|---|---|---|---|---|---|---|
| Adult | 1 | 2485 | NA | NA | NA | NA |
| Juvenile | 1 | 568 | NA | NA | NA | NA |
| Hatch | 1 | 26 | NA | NA | NA | NA |
| Juvenile & Adult | 2 | 390 | 106.74 | 3.65 | 2.89E-157 | 1.16E-156 |
| Hatch & Adult | 2 | 12 | 4.89 | 2.46 | 0.001 | 1.32E-03 |
| Hatch & Juvenile | 2 | 15 | 1.12 | 13.43 | 1.30E-14 | 2.60E-14 |
| Hatch & Juvenile & Adult | 3 | 9 | 0.21 | 42.88 | 3.75E-13 | 5.00E-13 |

**Supplemental table 8|** Number of sex-biased genes (SB) overlapping between stages (SB in № stages), showing a significant sex by developmental stage interaction (Sex by development interaction) in *T. californicum*.

| SB in № stages | SB | Sex by development interaction | % |
|---|---|---|---|
| **1** | 2272 | 811 | 35.7 |
| **2** | 390 | 253 | 64.9 |
| **3** | 9 | 3 | 33.3 |

**Supplemental table 9|** *Timema californicum* samples and their accession numbers

| Experiment Accession | Sample Accession | Run Accession | Library Name | Developmental stage and sex | Total Spots | Total Bases |
|---|---|---|---|---|---|---|
| SRX9531643 | SRS7738165 | SRR13084978 | Tcm_M_H_rep1 | hatchling_male | 44601473 | 9009497546 |
| SRX9531642 | SRS7738164 | SRR13084979 | Tcm_F_H_rep1 | hatchling_female | 44379870 | 8964733740 |
| SRX9531641 | SRS7738163 | SRR13084980 | Tcm_M_A_rep4 | adult_male | 37276088 | 7529769776 |
| SRX9531640 | SRS7738162 | SRR13084981 | Tcm_M_A_rep3 | adult_male | 35119725 | 7094184450 |
| SRX9531639 | SRS7738161 | SRR13084982 | Tcm_M_A_rep2 | adult_male | 39800843 | 8039770286 |
| SRX9531638 | SRS7738160 | SRR13084983 | Tcm_M_A_rep1 | adult_male | 36436465 | 7360165930 |
| SRX9531637 | SRS7738159 | SRR13084984 | Tcm_F_A_rep4 | adult_female | 39841828 | 8048049256 |
| SRX9531636 | SRS7738158 | SRR13084985 | Tcm_M_J_rep4 | juvenile_male | 19806861 | 4000985922 |
| SRX9531635 | SRS7738157 | SRR13084986 | Tcm_F_J_rep3 | juvenile_female | 43925792 | 8873009984 |
| SRX9531634 | SRS7738156 | SRR13084987 | Tcm_F_J_rep2 | juvenile_female | 43805442 | 8848699284 |
| SRX9531633 | SRS7738155 | SRR13084988 | Tcm_F_J_rep1 | juvenile_female | 47842363 | 9664157326 |

| | | | | | | |
|---|---|---|---|---|---|---|
| SRX9531632 | SRS7738154 | SRR13084989 | Tcm_F_A_rep3 | adult_female | 38345072 | 7745704544 |
| SRX9531631 | SRS7738153 | SRR13084990 | Tcm_M_J_rep3 | juvenile_male | 41828810 | 8449419620 |
| SRX9531630 | SRS7738152 | SRR13084991 | Tcm_M_J_rep2 | juvenile_male | 44470745 | 8983090490 |
| SRX9531629 | SRS7738151 | SRR13084992 | Tcm_M_J_rep1 | juvenile_male | 49269834 | 9952506468 |
| SRX9531628 | SRS7738150 | SRR13084993 | Tcm_F_H_rep4 | hatchling_female | 45158508 | 9122018616 |
| SRX9531627 | SRS7738149 | SRR13084994 | Tcm_M_H_rep5 | hatchling_male | 48849073 | 9867512746 |
| SRX9531626 | SRS7738148 | SRR13084995 | Tcm_F_H_rep3 | hatchling_female | 63932274 | 12914319348 |
| SRX9531625 | SRS7738147 | SRR13084996 | Tcm_M_H_rep4 | hatchling_male | 46547428 | 9402580456 |
| SRX9531624 | SRS7738146 | SRR13084997 | Tcm_F_H_rep2 | hatchling_female | 40703580 | 8222123160 |
| SRX9531623 | SRS7738145 | SRR13084998 | Tcm_M_H_rep3 | hatchling_male | 42144046 | 8513097292 |
| SRX9531622 | SRS7738144 | SRR13084999 | Tcm_M_H_rep2 | hatchling_male | 37358520 | 7546421040 |
| SRX9531621 | SRS7738143 | SRR13085000 | Tcm_F_A_rep2 | adult_female | 36838347 | 7441346094 |
| SRX9531620 | SRS7738142 | SRR13085001 | Tcm_F_A_rep1 | adult_female | 38243875 | 7725262750 |

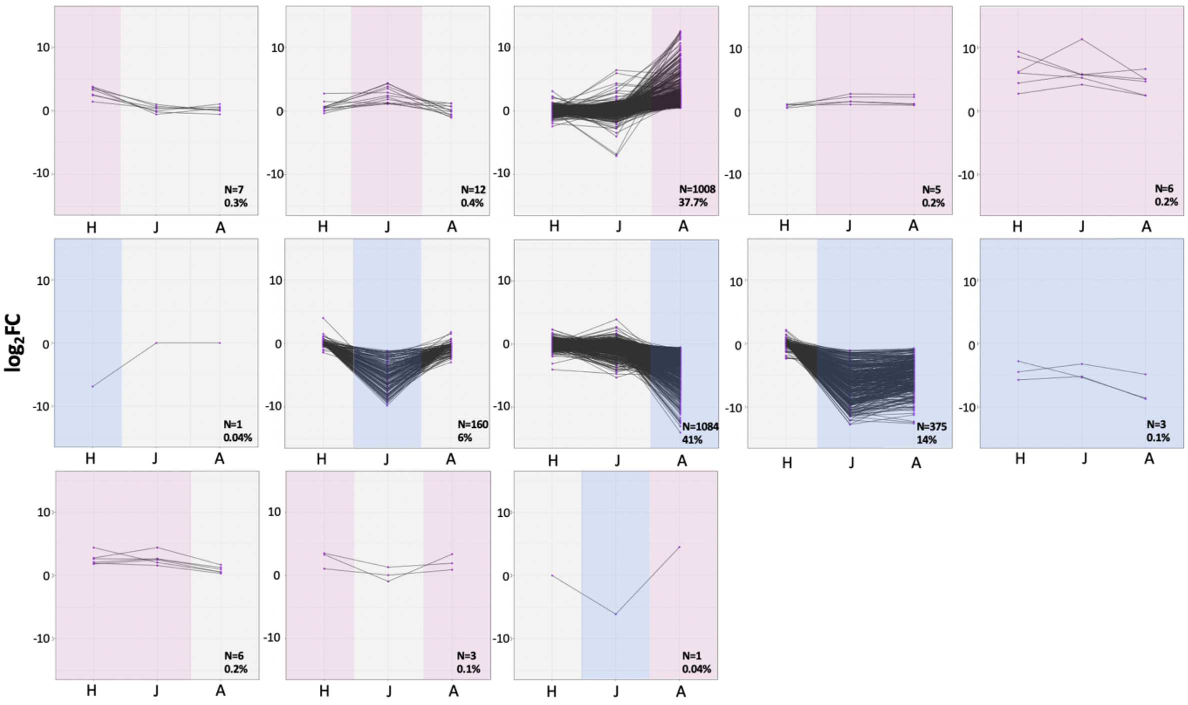


**Supplemental figure 1|** Categories of sex-biased genes in *T. californicum* ("H"- hatchling, "J"- juvenile, "A"- adult). Blue background stands for male-biased state of a gene, pink background stands for female-biased state of a gene, and gray background stands for un-biased state of a gene. For each category the number of genes (N) and their percentage is marked at the bottom right corner.

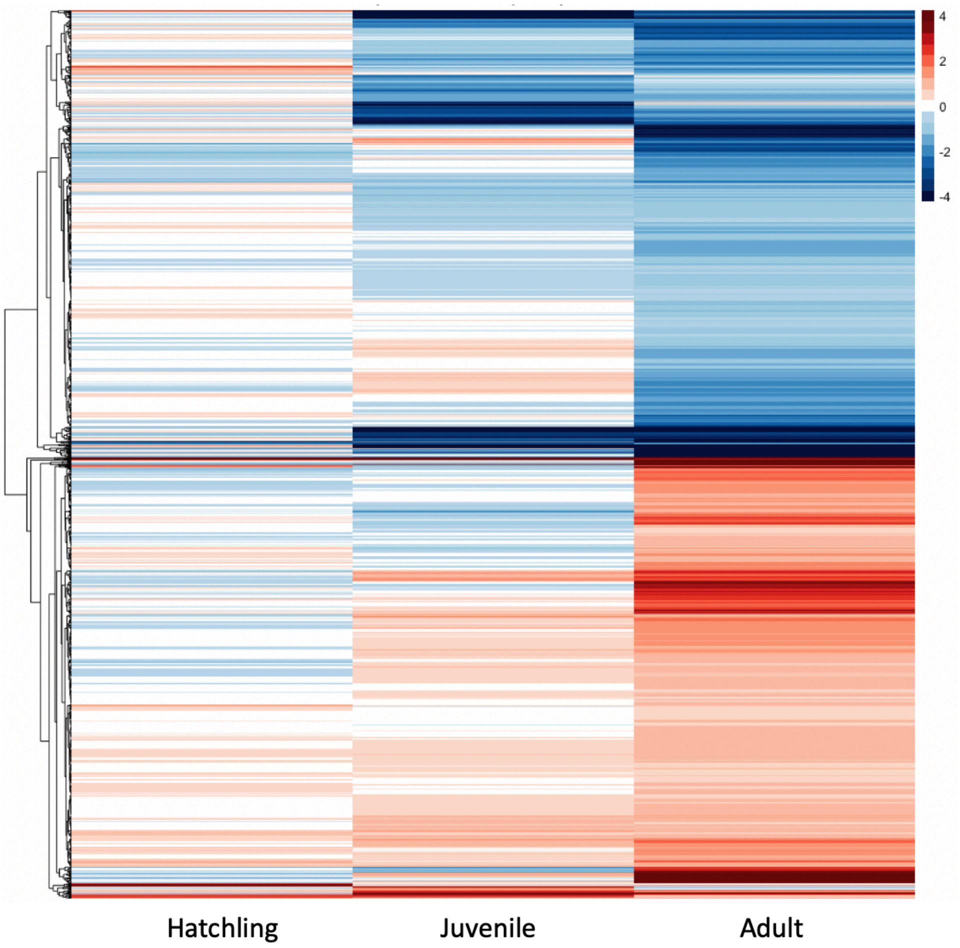


**Supplemental figure 2|** Heat-map showing the $log_2FC$ over three developmental stages of 1887 genes that were significantly sex-biased in at least one stage. Note that a subset of genes (n=784) was removed due to low expression in some of the stages. Genes in red are female-biased, blue are male-biased.

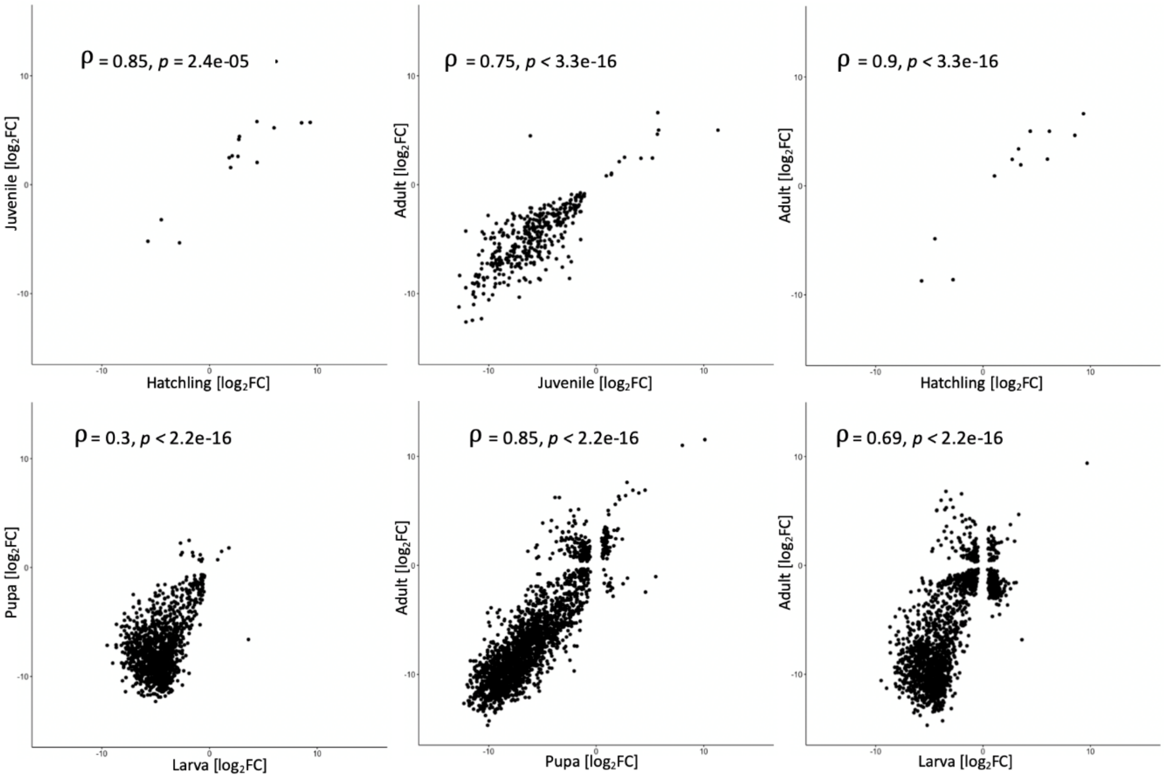


**Supplemental figure 3 |** Spearman's correlations of sex-biased gene expression [$log_2FC$] between developmental stages. Top panels: *T. californicum*; bottom panels: *D. melanogaster.* Spearman's correlation coefficients and adjusted p values are shown in each panel.

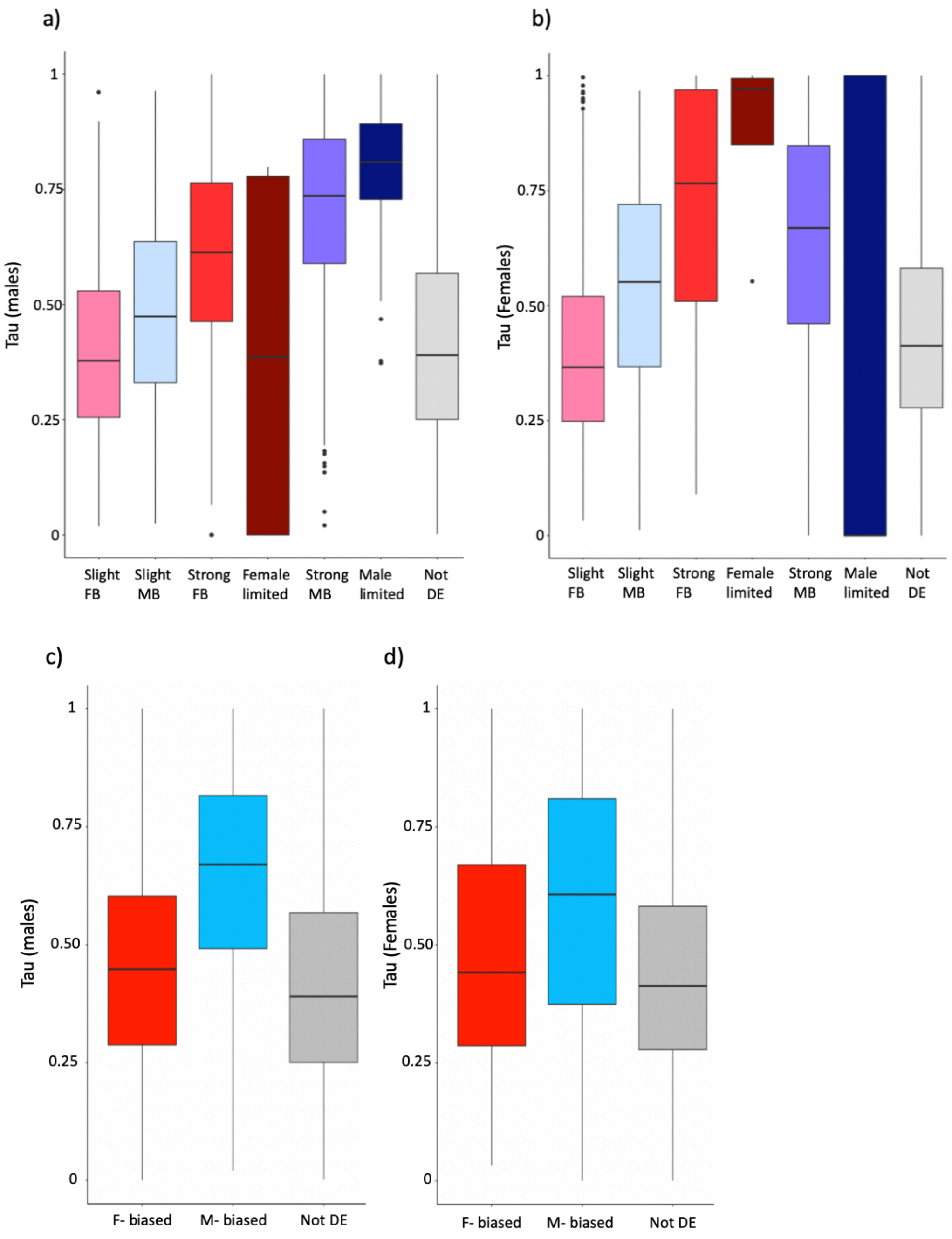


**Supplemental figure 4 |** Tau index of gene expression in males (a, c) and females (b, d), for each sex-bias category; slight FB, slight MB, strong FB, strong MB, Female-limited, Male limited, Not DE (a,b), and per broader sex-bias categories; F-biased, M-biased and Not DE. Tau ranges from zero (uniformly expressed over development) to one (gene expressed in only one stage). Gene categories are depicted with different colors; boxplots represent the median, lower and upper quartiles, and whiskers the minimum and maximum values (in the limit of 1.5x interquartile range).

**a)** Hatchlings

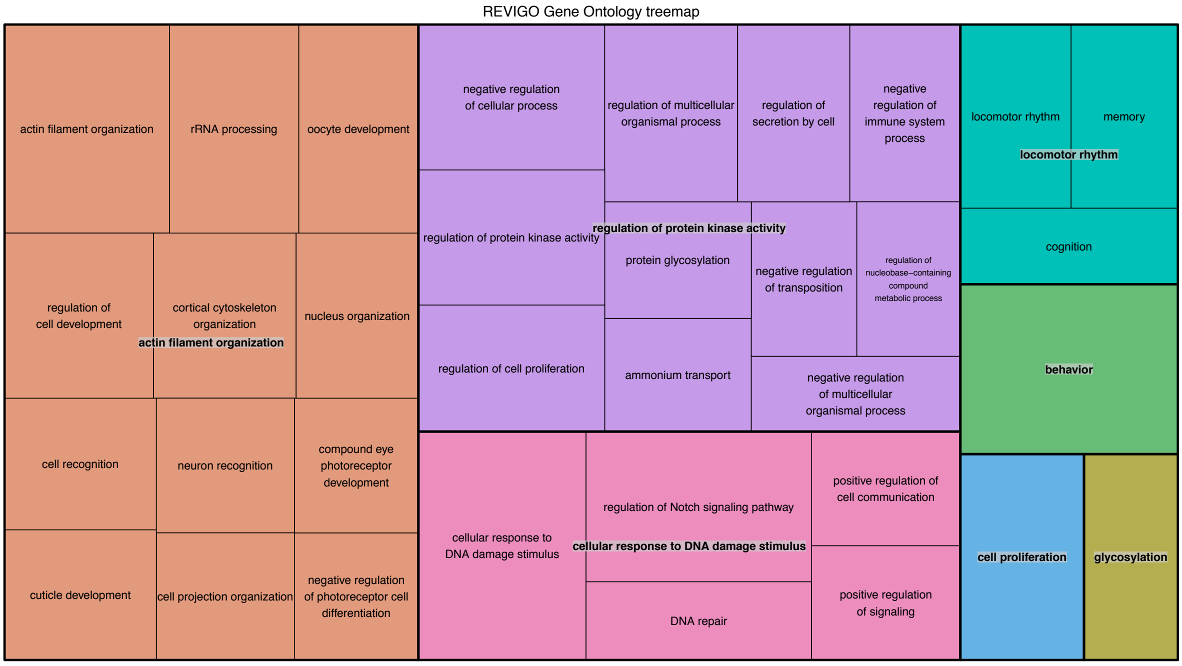


**b)** Juveniles

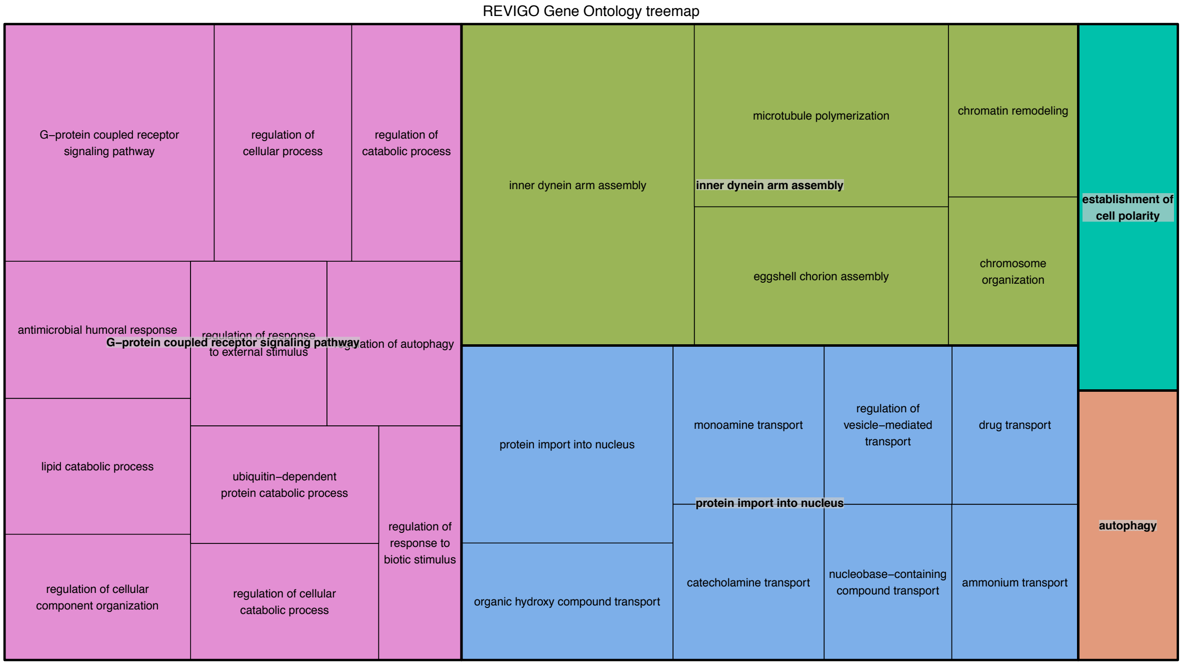

**c)** Adults

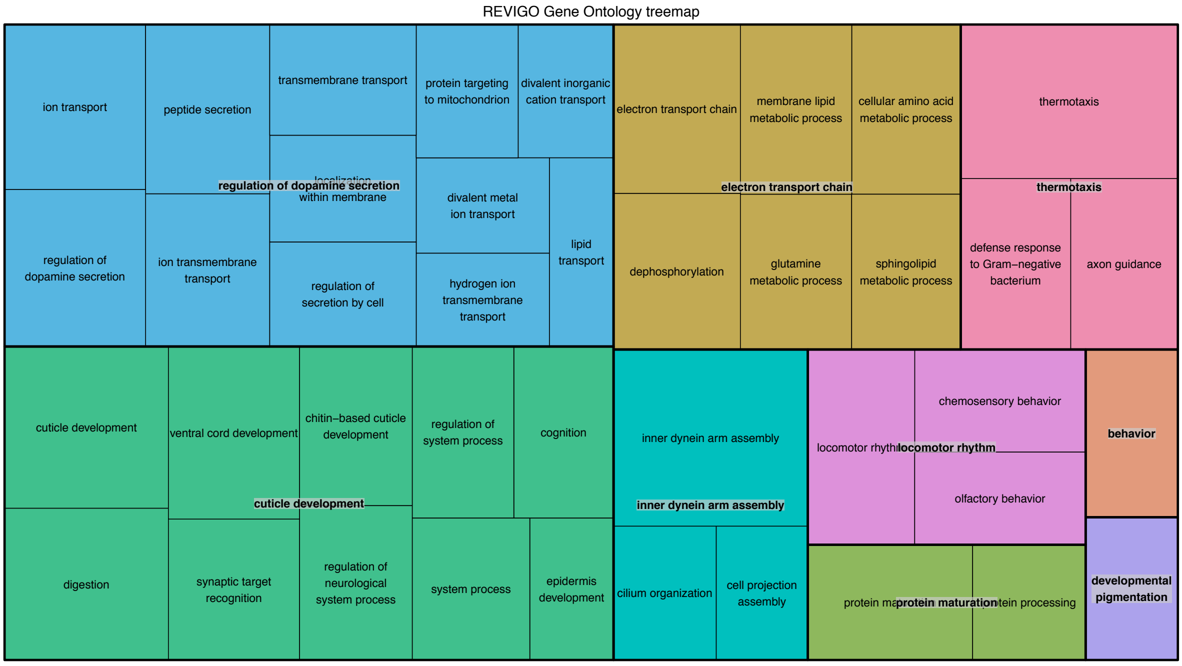


**Supplemental figure 5 |** Summary of significant GO terms by Revigo (Supek et al., 2011) for genes that are sex-biased in a) hatchlings, b) juveniles and c) adults.
